# Ribosomal protein S1 plays a critical role in horizontal gene transfer by mediating the expression of foreign mRNAs

**DOI:** 10.1101/2021.10.13.464283

**Authors:** Luc Roberts, Hans-Joachim Wieden

## Abstract

The emergence of multi-antibiotic resistant bacteria is one of the largest threats to global heath. This rise is due to the genomic plasticity of bacteria, allowing rapid acquisition of antibiotic resistance through the uptake of foreign DNA (i.e. horizontal gene transfer, HGT). This genomic plasticity is not limited to DNA from bacteria, highly divergent (trans-kingdom) mRNA have been reported to drive translation in *E. coli*. Trans-kingdom activity has been attributed to mRNA tertiary structure suggesting the bacterial translation machinery bottle-necks HGT, restricting the expression of foreign DNA. However, here we show that tertiary structure is not responsible for ribosome recruitment and that the translation efficiency is dependent on ribosomal protein S1 and an A-rich Shine-Dalgarno-like element. The S1-facilitated ability of ribosomes to identify and exploit A-rich sequences in foreign RNA highlights the important role that S1 plays in horizontal gene transfer, the robustness of canonical prokaryotic translation, and bacterial evolution.

## INTRODUCTION

Horizontal gene transfer (HGT) is the transmission of genetic information between two organisms other than by reproduction (i.e. parent to offspring). HGT plays a major role in bacterial evolution, highlighted by the fact that it is the primary mechanism by which antibiotic resistance genes are shared ^1-3^. One mechanism by which bacteria transfer genes horizontally is through active uptake of DNA from their environment, a process known as natural transformation ^4^. Natural transformation is advantageous from an evolutionary standpoint as it allows bacteria to increase their genetic variability thereby exploring a larger potential fitness landscape. However, these advantages rely on the ability of the bacteria to transcribe and subsequently translate the newly acquired genetic information into functional gene products. Foreign DNA transcription has been observed in *E. coli* and appears to inversely correlate with phylogenetic distance from the host as roughly 50% of the *H. influenza* genes are transcribed compared to relatively few human genes ^5^. Additionally, a fragment of the human long interspersed nuclear element L1 has been found in a *Neisseria gonorrhoeae* genome and is actively transcribed ^6^, suggesting that bacteria are capable of utilizing foreign DNA from evolutionary distant sources. While previous work has revealed that the sporadic transcription of foreign DNA is due to a combination of specific and non-specific mechanisms inherent to bacteria ^5,7^, they have failed to elucidate whether the resulting “foreign” mRNAs are able to compete with native bacterial mRNAs and ultimately contribute to the proteome of the recipient cell (Figure 1). The contributing factors within the cell that govern the translation of horizontally acquired genes are particularly interesting both, from an evolutionary point of view as well as with respect to the mechanisms that contribute to the emergence of antimicrobial resistance, as foreign mRNA (derived from an evolutionarily distant DNA) will not have a translation initiation region that is compatible with the bacterial translation initiation machinery. Does the translation machinery serve as a filter for foreign genetic information to avoid detrimental effects to the cell caused by incompatible / toxic gene products? Or is translation fundamentally promiscuous suggesting that HGT is not limited to DNA from closely related (evolutionarily) species? In support of the former, recently an internal ribosome entry site (IRES) from the insect pathogen *Plautia Stali* intestine virus (PSIV) was reported to drive structure-based translation in *E. coli*. IRESs are RNA elements capable of recruiting ribosomes and initiating translation on an internal portion of an mRNA ^8,9^. The intergenic region (IGR) IRES of the Dicistroviridae virus family is unique as it does not require any initiation factors and initiates on a non-AUG start codon ^10-12^. This non-canonical activity relies on a triple pseudoknot (PK) tertiary structure, which binds to the ribosomal subunits using this structural element mimicking a canonical tRNA-mRNA duplex ^13-20^. However, the reported prokaryotic IRES activity differs from that in the eukaryotic context. In *E. coli* the AUG start codon is essential for translation, additionally disrupting PK structures known to be essential for translation initiation in Eukaryotes has minimal effects on translation efficiency (Colussi et al., 2015). As a consequence, the proposed mechanism for prokaryotic IRES activity is a “hybrid” of the previously described eukaryotic IRES activity and the canonical Shine-Dalgarno (SD) initiation mechanisms whereby bacterial ribosomes transiently interact (in a structure dependent manner) with the IRES before repositioning to a downstream SD-like sequence (Colussi et al., 2015). This suggests some foreign mRNAs can indeed be specifically translated in bacteria by exploiting the evolutionary conserved structural features of the ribosomal core ^21,22^. However, this would significantly limit the “foreign” transcripts that can be expressed to those that have for a particular reason in their native context evolved to exploit conserved features of the translation machinery. Therefore, we hypothesized that in order to optimally utilize horizontally acquired DNA the bacterial translation machinery should be at some level fundamentally promiscuous.

**Figure 1.**
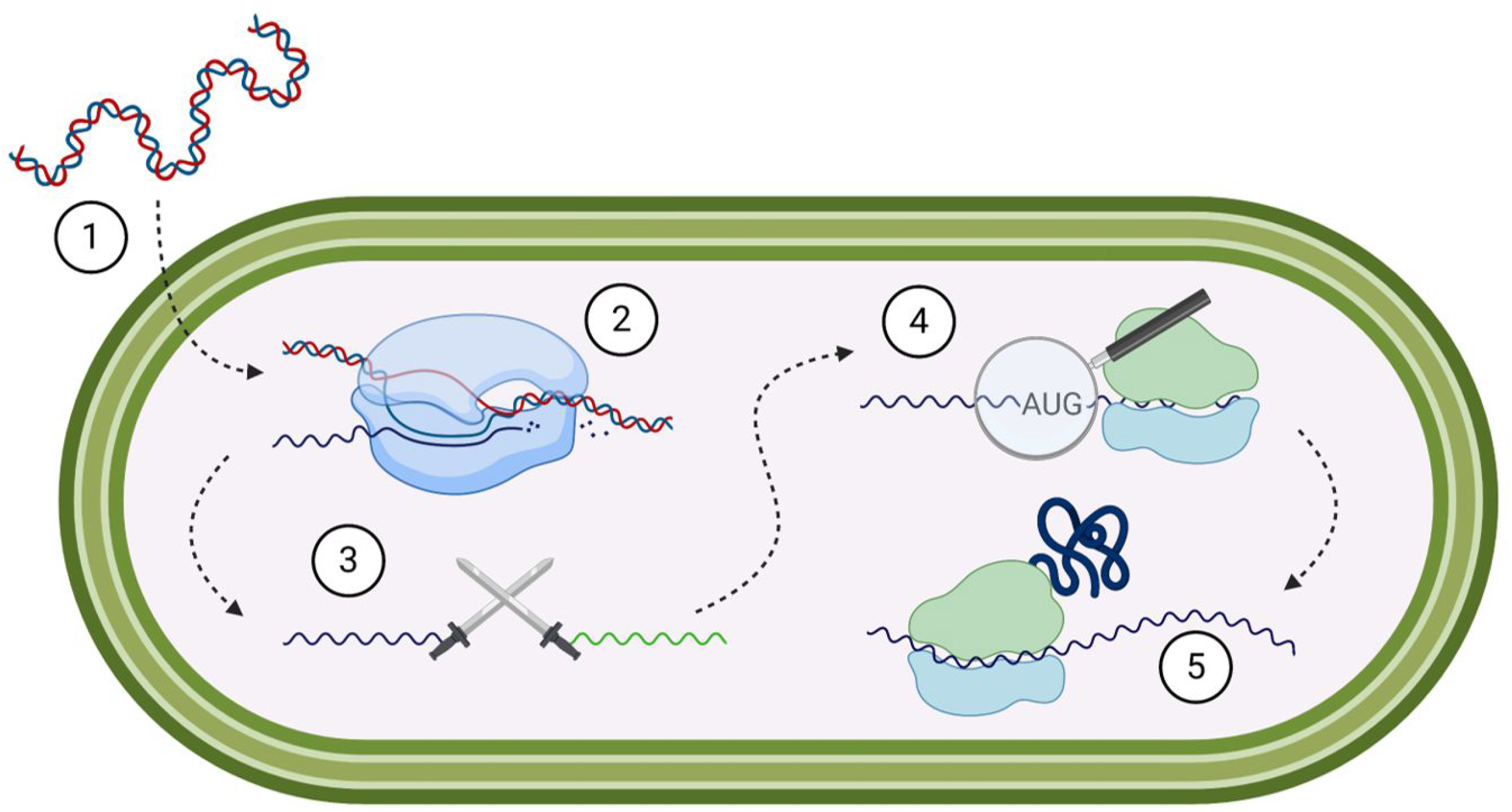
Foreign gene expression in *E. coli*. (1) Foreign DNA taken up into the cell. (2) Spurious transcription generates foreign mRNA which (3) competes with native mRNAs (green) for ribosome binding. (4) Upon locating a start codon (5) ribosomes translate the gene generating a foreign protein.

To determine the biological underpinnings of foreign DNA expression, as well as the minimal/mechanistic requirements for effective foreign mRNA translation, we performed the first detailed single cell *in vivo* quantitative characterization of foreign mRNA (viral IRES) driven translation in *E. coli*. Our results demonstrate that foreign mRNA translation correlates positively with AT-rich regions, is mediated by ribosomal protein S1 and that a SD-like sequence upstream of the start codon is required for efficient translation. This supports the notion that the bacterial translation machinery is not inherently biased to translate specific foreign mRNAs but rather seems to allow for effective translation of any mRNA, who’s structure can be resolved by ribosomal protein S1 and where a SD-like sequence upstream of a start codon can be identified. This clearly highlights the critical role that S1 plays for the strategy of unbiased sampling of foreign mRNAs (from viruses, eukaryotes, other prokaryotes, etc.) in bacteria consistent with maximizing access to potentially useful (antimicrobial resistance) DNA acquired via horizontal gene transfer.

## RESULTS

To investigate the question if trans-kingdom expression of foreign DNA is able to compete with endogenous translation signals in actively translating bacteria and ultimately contribute to the proteome of the recipient cell (Figure 1), we developed a real-time fluorescence-based single-cell translation assay. We opted to measure individual live cell fluorescence by flow cytometry to avoid potential averaging of distinct *E. coli* populations and to provide information regarding the population wide behaviour of the translation initiation event. To benchmark our reporter system, we selected three ribosome binding sites (RBSs) (Strong B0034, Medium B0032, and Weak B0033) from the registry of standard biological parts (http://parts.igem.org) ^23^, as well as a “dead” RBS (the reverse complement of B0034), to drive the expression of superfolder green fluorescent protein (sfGFP) and analysed their translation efficiency (TE) using flow cytometry (Figure S1). The obtained flow cytometry measurements correlate nearly perfectly with the predicted expression strength (ρ = 0.99) using the ribosome binding site (RBS) calculator, demonstrating the sensitivity and wide range over which our single-cell assay can accurately report translation efficiency *in vivo* without interference by endogenous expression even at low expression levels (Figure 2A)^24^. By benchmarking foreign mRNAs against well-characterized SDs we are able to accurately measure their translation efficiency allowing for the first time a direct comparison of foreign mRNA translation to the canonical system, assessing their ability to compete with native mRNAs *in vivo*.

**Figure 2.**
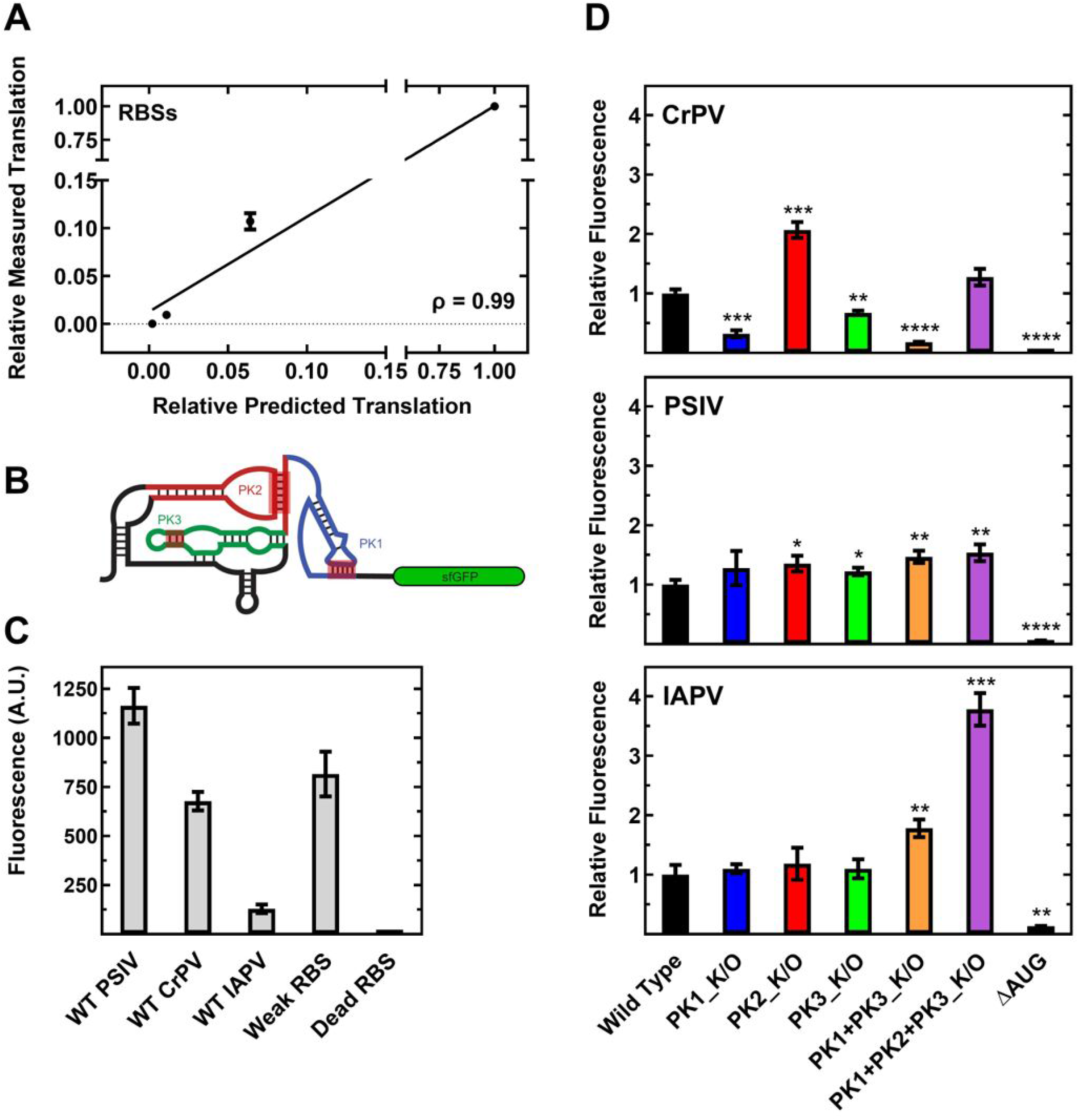
IGR IRES translation efficiency is independent of pseudoknot structure. (A) Correlation of predicted and measured translation efficiency of standardized RBSs. Translation efficiency predicted using the Salis lab RBS calculator, translation efficiency measured by flow cytometry. Mean values of three biological replicates are plotted. (B) Cartoon representation of the monocistronic fluorescent reporter construct including the secondary structure of the IGR IRESs. Pseudoknots (PK) 1 (blue), 2 (red), and 3 (green) are indicated. Location of pseudoknot mutations are highlighted in red, the exact sequences are summarized in Table S4. (C) Translation efficiencies measured by flow cytometry of the WT IGR IRESs compared to standardized RBSs (D) Translation efficiency measured by flow cytometry, mean values of three biological replicates are plotted relative to the respective WT IRES. For all panels error bars indicate one standard deviation. Constructs with statistically significant differences from WT are indicated (* = P < 0.05, ** = P < 0.01, *** = P < 0.001, **** = P < 0.0001).

### 1) The IGR IRESs have translational efficiencies comparable to a weak RBS in vivo

In order to assess and quantify the ability of a foreign translation initiation element to drive cross-kingdom gene expression at the translation level we decided to utilize IGR IRESs. We initially selected the cricket paralysis virus (CrPV) IRES as it is arguably the best characterized of the IGR IRESs. To accurately mimic the expression of a foreign sequence we included 18nts of viral coding sequence downstream of the IRES to keep the initiation element in its native context and our reporter system (Figure 2B) consistent with previous studies ^12,22^. Finally, we opted for a monocistronic IRES construct to avoid potential translational coupling, as downstream translation and the intergenic RNA structure can be influenced by upstream translation ^25,26^. The obtained live-cell fluorescence data revealed that within actively translating ribosomes the translation efficiency (TE) of the wild type (WT) CrPV construct is roughly equivalent to the TE observed for a weak bacterial RBS (Figure 2C). A low translation level is consistent with the observation that in eukaryotes IRES translation can be outcompeted by canonical eukaryotic translation ^27^. In agreement with previous primarily qualitative studies deletion of the sfGFP start codon abolishes translation (Figure 2D, top panel) ^22^, suggesting an initiator tRNA dependent transition from initiation to elongation *in vivo*, similar to the canonical process in bacteria.

### 2) Disruption of PK structure does not perturb IRES activity in vivo

To investigate the detailed mechanistic role that the structured region of the IRES plays in its translational efficiency, we systematically disrupted the conserved pseudoknots (PK) (Figure 2B) present in the CrPV IRES. The PKs are critical for translation initiation in eukaryotes and it has been previously demonstrated that altering the sequence of the IRES to disrupt the Watson-Crick-Franklin base pairs affects only the PK structure leaving the rest of the IRES intact ^12^. In line with this, disrupting PK1 (PK1_K/O) and PK3 (PK3_K/O) decreases the translation efficiency 70% and 30% respectively and when combined (PK1+PK3_K/O) decrease TE 80% (Figure 2D). Surprisingly, disruption of PK2 (PK2_K/O) increases translation efficiency 2-fold, and when combined with the other PK mutations (PK1+PK2+PK3_K/O), restores activity to WT level (Figure 2D). The observation that disrupting all pseudoknots (PK1+PK2+PK3_K/O) in the CrPV IRES does not change the translation efficiency suggest that its tertiary structure is not required for the observed cross-kingdom expression activity of the CrPV IRES. Complete removal of PK elements further confirms this as deletion of the highly structured PK2 and PK3 (ΔPK2/3) results in a 4-fold increase in TE while deletion of the relatively less structured PK1 (ΔPK1) abolishes translation (Figure S2A).

Given that these results are inconsistent with the proposed structure-driven translation initiation mechanism reported for the PSIV IRES ^22^ and to rule out any effects specific to the CrPV IRES, we decided to analyse the PSIV IRES as well as the Israeli Acute Paralysis Virus (IAPV) IRESs. Interestingly, although structurally similar to CrPV the WT PSIV IRES has a 2-fold greater translation efficiency than the weak RBS, while the WT IAPV IRES translation efficiency is only 15% of the weak RBS (Figure 2C). Disruption of PK1, PK2, PK3, and combinations of these mutations have no negative effects on IAPV or PSIV translation, and like CrPV, several constructs exhibit increased translation efficiency (∼1.2 – 4-fold) compared to their WT counterparts (Figure 2D, Table 1). To assess the inherent translational activity of the IGR IRESs primary sequences without their tertiary structure we randomized the CrPV and IAPV primary sequences (while maintaining nucleotide composition). Interestingly, the randomized CrPV and IAPV translation efficiencies are ∼3 and ∼7-fold greater than their WT counterparts (Figure S2 B and C), this brings CrPV translation efficiency 3-fold over the weak RBS and elevates IAPV to the level of the weak RBS. Together our results indicate that the tertiary structure of the IRES is not responsible for, but instead is inhibitory to, efficient translation. Therefore, we hypothesized that the IRESs are being treated as large structured 5’ UTRs and translated via the canonical processes of translation machinery that deal with the expression of structured mRNA rather than through molecular mimicry (as is the case in Eukaryotes). To determine if this IGR IRES translational activity is indeed due to canonical translation, we performed a detailed characterization of the PSIV IRES (Figure 3A) driven translation.

**Table 1.** Relative *in vivo* translation efficiencies of IGR IRES constructs measured by flow cytometry. Mean values of three biological replicates are plotted; error bars indicate one standard deviation.

|  | **PSIV** | **IAPV** | **CrPV** |
| --- | --- | --- | --- |
| **Wild Type** | 1.00 ± 0.08 | 1.00 ± 0.17 | 1.00 ± 0.07 |
| **PK1_K/O** | 1.28 ± 0.29 | 1.10 ± 0.08 | 0.32 ± 0.06 |
| **PK2_K/O** | 1.35 ± 0.13 | 1.18 ± 0.03 | 2.07 ± 0.13 |
| **PK3_K/O** | 1.22 ± 0.06 | 1.10 ± 0.16 | 0.67 ± 0.04 |
| **PK1+PK3_K/O** | 1.47 ± 0.11 | 1.78 ± 0.15 | 0.18 ± 0.01 |
| **PK1+PK2+PK3_K/O** | 1.54 ± 0.14 | 3.78 ± 0.28 | 1.29 ± 0.14 |
| **SD-like_K/O** | 0.04 ± 0.01 | - | - |
| **sfGFP ΔAUG** | 0.06 ± 0.01 | 0.14 ± 0.01 | 0.04 ± 0.01 |

**Figure 3.**
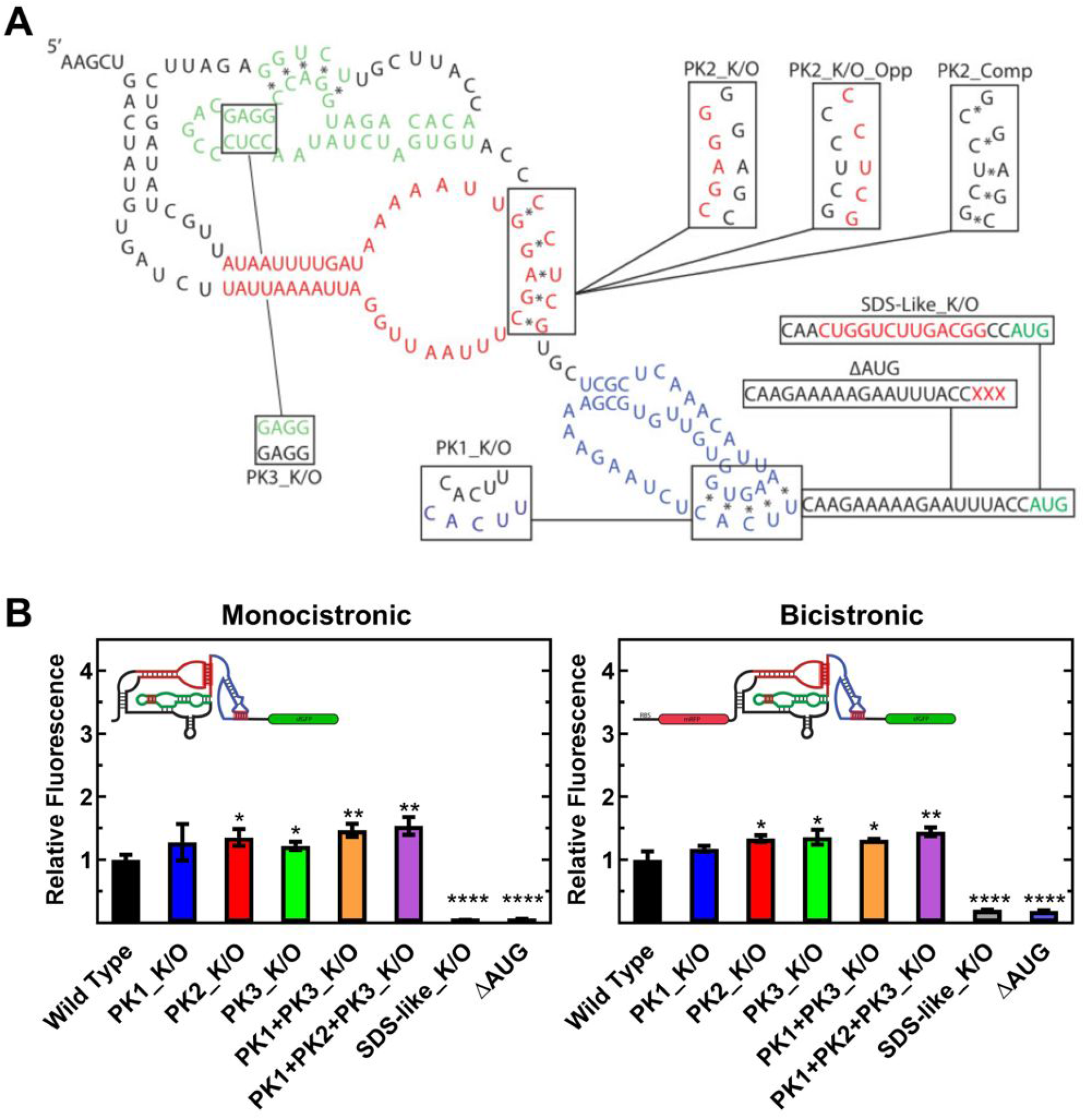
PSIV IGR IRES translation efficiency is independent of its position in the mRNA. (A) Secondary structure of the PSIV IRES with nucleotide substitutions for annotated mutations shown in red. (B) Translation efficiency measured by flow cytometry, mean values of three biological replicates are plotted relative to the respective WT IRES; error bars indicate one standard deviation. Constructs with statistically significant differences from WT are indicated (* = P < 0.05, ** = P < 0.01, *** = P < 0.001, **** = P < 0.0001).

### 3) IRES location has no effect on translation initiation mechanism selection

In their natural context the IGR IRESs initiate translation in the intergenic region of a bicistronic viral genome ^9,28^. While canonical Shine-Dalgarno based translation, according to the definition, could be considered “IRES” translation, it is possible that the structure of the IRES will be more efficient at promoting initiation if located between two genes as opposed to the monocistronic reporter we utilized (Figure 2B). To test this, we designed a bicistronic reporter construct whereby an upstream mRFP is translated via the weak RBS (the most similar to the WT PSIV IRES expression level) and the PSIV IRES drives the translation of sfGFP downstream (Figure S3). This reporter design is often used to validate IRES activity in Eukaryotes and allows calculation of the mRFP/sfGFP ratio to control for intrinsic cellular noise. Subsequent analysis using flow cytometry revealed that the relative mRFP/sfGFP expression levels of the WT PSIV and PK variants are identical to the monocistronic expression levels (Figure 3B). These results demonstrate that the location of the IRES (5’ or internal) is not biasing against a structure-based initiation mechanism and reaffirms that the structure of the IRES is not essential for efficient translation.

### 4) IRES translational efficiency is independent of bacterial growth phase

In our initial experiments we measured fluorescence at a single time point three hours post induction, shortly after entry into the stationary growth phase. At this point expression of endogenous mRNAs is reduced which might increase the translational efficiency of the IRES by reducing competition with these mRNAs. It is possible that during times of increased competition for ribosomes (rapid growth) the structure of the IRES provides a kinetic advantage to the mRNA by transiently interacting with ribosomes in a structure dependent manner. In order to investigate if the structure of the IRES provides cross-kingdom activity during rapid growth we analyzed IRES translation efficiency during different growth phases. To this end we performed an in-depth time course analyses of the WT PSIV and PK2_K/O PSIV IRES constructs. As our fluorescence-based assay is able to accurately quantify the amount of sfGFP present *in vivo* (Figure S4) and is not limited by sfGFP degradation (Figure S5) or sfGFP maturation, we were able, for the first time, to investigate the ability of a foreign mRNA to compete with endogenous mRNAs for translation over several growth phases (lag, exponential, and early stationary, Figure S6). In agreement with our initial data, PK2_K/O reaches a higher final fluorescence (Figure 4A) and the rate of sfGFP production during the exponential phase is 2-times greater for PK2_K/O (41.1 ± 1.4 min^-1^) than for WT (19.9 ± 0.6 min^-1^) (Figure 4B). This demonstrates that foreign mRNA translation is robust and can successfully compete with endogenous mRNA expression under multiple growth conditions, providing further evidence that for their translation the IRES structure is not required and might even limit TE.

**Figure 4.**
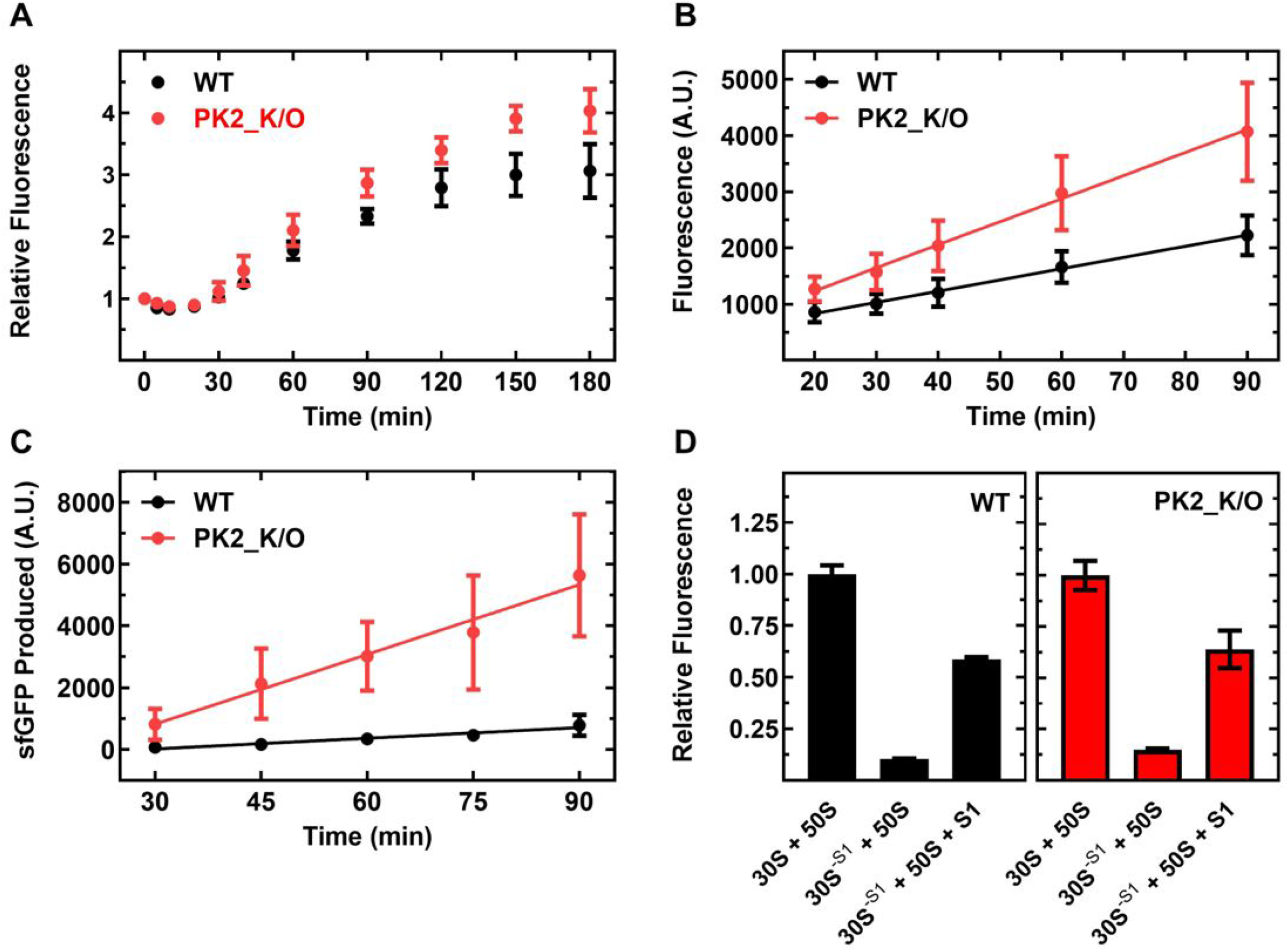
PSIV IGR IRES translation efficiency is consistent over multiple growth phases. (A) Relative fluorescence *in vivo* time course of E. coli containing PSIV IGR IRES constructs as measured by flow cytometry. (B) The linear portion of sfGFP expression in panel A (C) sfGFP production *in vitro* time course using the PSIV IGR IRES constructs and the PURExpress® system (D) Relative fluorescence of PSIV IGR IRES constructs and the PURExpress® Δ ribosome system and ribosomes with and without ribosomal protein S1. Mean values of three biological replicates are plotted; error bars indicate one standard deviation.

### 5) Disruption of IRES PK structure affects mRNA stability in vivo

Although the PK structure seems not to be important for the TE in bacteria some of the PK variants show decreased expression consistent with a structure-based initiation mechanism. Disruption of the PK structure, while only affecting local RNA structure (^12^, could be affecting the stability of the mRNA *in vivo* and therefore as a consequence reduce the translational efficiency of the IRES variants, providing an explanation for the observed effect. To assess mRNA accumulation and stability at different growth phases, we measured the total mRNA levels via qRT-PCR on samples collected during the in-depth time course analysis. Interestingly, while mRNA levels were similar in the early stationary phase (Figure S7A) WT mRNA accumulated faster (∼ 2x) than PK2_K/O mRNA, supporting the notion that disruption of the PK2 (PK2_K/O) decreases mRNA stability (Figure S7B). This result also suggests that the PK2_K/O has an even higher relative translational efficiency than reported by our fluorescence data when factoring in its lower abundance due to the reduced stability of the mRNA.

### 6) IRES translation may be due to ribosome standby sites

We noticed a trend in our data that PK variants with increased TE almost always contained substitutions that introduced new single stranded (or previously base-paired) AG-rich “SD-like” sequences upstream of the start codon, which have been shown previously to promote initiation ^29^. These single stranded SD-like sequences could affect the translational efficiency of the IGR IRESs by acting as ribosome standby sites, S1 binding sites, or simply by facilitating the breathing of neighbouring RNA structures ^30,31^. As a simple check for this we introduced a compensatory mutation that re-establishes PK2 structure (PK2_Comp) and that returns TE to WT level (Figure S8). However, the compensatory mutation alone PK2 (PK2_Opp), while disrupting PK structure, does not introduce a single stranded SD-like sequence (Figure 3A) and has no effect on IRES translation efficiency (Figure S8). This suggest (at least for this particular variant) that not the disruption of PK structure but rather the presence of a single stranded AG-rich (SD-like) sequence upstream of the start codon is important for increased translation efficiency.

### 7) Ribosomal Protein S1 is required for efficient IRES translation

Our *in vivo* assays indicate that the structure of the IRES is non-essential and in fact even limit translational efficiency, suggesting that additional factors such as RNA helicases to resolve the structure of the IRES might be required for efficient translation. To determine if cellular factors or the ribosome itself are responsible for this we measured the rate of IRES-mediated sfGFP production *in vitro* using the highly purified and reconstituted PURExpress® system ^32^. The respective sfGFP synthesis time courses mirror our *in vivo* data, as the rate of sfGFP production is 6-times greater for PK2_K/O (75.3 ± 25.3 min-1) than for WT (11.5 ± 4.7 min-1) (Figure 4C). This supports the notion that the component responsible for the observed effect is present in the recombinant, purified and reconstituted coupled transcription and translation system, and also backs our earlier observation that the *in vivo* translation efficiency of the PSIV IRES PK2_K/O might be limited by decreased *in vivo* stability of the respective mRNA (Figure S7) as the contributing nuclease are not present in the PUREexpress® system. Within the PURExpress® system ribosomal protein S1 is the most likely candidate to resolve the IRES structure ^33^ as S1 binds to single stranded A-rich sequences and possesses helicase activity essential for efficient canonical translation of mRNAs with structured 5’ UTRs in *E. coli* ^31^. If the IRESs are indeed being treated as large structured 5’ UTRs by the bacterial translation machinery, their translational efficiency will be reliant on S1. However, if specific interactions of the triple PK structure of the IRES with the ribosome are responsible for its’ translational efficiency, the absence of S1 should have little to no effect. To probe this, we monitored IRES translation efficiency, using the PURExpress® delta ribosome kit supplemented with either ribosomes (30S + 50S) or ribosomes lacking S1 (30S^-S1^ + 50S). Both the WT and PK2_K/O translation efficiencies were decreased ∼90% when S1 was not present (Figure 4D), likely due to the ribosome no longer being able to efficiently bind and unwind the highly structured mRNA. To ensure this is a S1 specific effect and not due to the treatment of the ribosomes during S1 removal we supplemented stoichiometric amounts of recombinant S1 (30S^-S1^ + 50S + S1) resulting in a 60% recovery in activity for both constructs (Figure 4D), demonstrating that S1 is indeed responsible for translation of these structured mRNAs.

### 8) IRES translation efficiency is dependent on a downstream SD-like sequence

If the IGR IRESs are being translated via the canonical translation initiation mechanism then the 30S ribosome will require a SD-like sequence adjacent to the sfGFP start codon for efficient translation ^29^. Interestingly, such a SD-like sequence has been identified in the viral coding region downstream of the PSIV IRES tertiary structure (^22^ and Figure 3A). Altering this sequence to its’ reverse complement (leaving the IRES structure intact) abolishes translation in both the mono- and bicistronic reporters (Figure 3B) further demonstrating that translation is proceeding via the canonical translation mechanism. Together with the S1 data this points to initiation being the determining factor in foreign translation efficiency.

### 9) Pseudoknot mutations do not perturb IGR IRES binding to the prokaryotic ribosome

While IRES PK structure is not responsible for the observed translation activity in prokaryotes, it is still possible that the IGR IRES is able to transiently interact with prokaryotic ribosomes in a structure specific manner that may be overshadowed by the canonical translation activity. In such a model disrupting the PKs structures will result in a reduced affinity of the IRES for the ribosome (as it is the case for Eukaryotic ribosome binding ^12^, Table S1). To investigate if the tertiary structure of the IGR IRES is responsible for prokaryotic ribosome binding, we determined the equilibrium binding constants (*K*_*D*_) for the CrPV and IAPV IRES-ribosome complexes using nitrocellulose filter binding.

WT CrPV and IAPV IRESs bind the 30S subunit and the 70S ribosome with comparable affinities of ∼100nM (Table 2). As expected, disrupting any of the PKs, individually or in combinations, has little to no effect on the affinity of either the CrPV or IAPV IGR IRESs for the 30S ribosomal subunit or the 70S ribosome (Table 2). This indicates that in contrast to the eukaryotic system (Table S1) the IGR IRESs bind to the prokaryotic ribosome independently of their tertiary structure ^12^. This is consistent with the observation that prokaryotes do not have an equivalent to eS25, the eukaryotic ribosomal protein shown to be critical for ribosome binding and IRES activity ^19,34,35^. To ensure that the use of nitrocellulose filtration to determine the affinity for the respective mRNAs is not biasing against a transient population of structurally bound IRESs we also measured the affinity for the 30S using equilibrium fluorescence titrations with a fluorescently labeled WT CrPV IRES, which resulted in a nearly identical affinity of ∼70nM (Figure S9). Finally, for comparison we tested two native structured 5’ UTRs (*rpsO* and *sodB*) and found that their affinities to the 30S and 70S ribosome are on the same order of magnitude as the IRESs (Table S2 and S3), further demonstrating the ability of foreign mRNAs to efficiently compete with native mRNAs for ribosome binding.

**Table 2.** Dissociation constants (K_D_) for CrPV and IAPV IGR IRES variants to 30S and 70S ribosomes as measured by nitrocellulose filter binding. Mean values of three biological replicates are plotted; error bars indicate one standard deviation.

| | | $K_D$ (nM) | $K_D$ (nM) |
| --- | --- | --- | --- |
| Construct | Ribosome | CrPV | IAPV |
| WT | 70S | 69 ± 7 | 50 ± 9 |
| PK1_K/O | 70S | 92 ± 9 | 67 ± 3 |
| PK2_K/O | 70S | 100 ± 13 | 59 ± 11 |
| PK3_K/O | 70S | 90 ± 15 | 59 ± 4 |
| PK1+PK3_K/O | 70S | 78 ± 10 | 65 ± 4 |
| PK1+PK2+PK3_K/O | 70S | 79 ± 10 | 72 ± 14 |
| WT | 30S | 110 ± 23 | 93 ± 7 |
| PK1_K/O | 30S | 55 ± 11 | 87 ± 14 |
| PK2_K/O | 30S | 74 ± 13 | 81 ± 25 |
| PK3_K/O | 30S | 48 ± 5 | 103 ± 42 |
| PK1+PK3_K/O | 30S | 52 ± 12 | 80 ± 28 |
| PK1+PK2+PK3_K/O | 30S | 48 ± 7 | 73 ± 14 |

## DISCUSSION

The ability of bacteria to exchange genes horizontally allows them to rapidly alter their genomic makeup, providing a critical fitness advantage when faced with environmental stresses such as antibiotics. To be effective, a system of active foreign DNA uptake must be paired with a cellular machinery that ensures that the newly acquired genes are transcribed and successfully translated to enable the corresponding fitness benefit. Here we sought to investigate the mechanisms utilized by bacteria to translate mRNAs derived from evolutionary distant foreign DNA, by performing an in-depth characterization of IGR IRES mediated translation in *E. coli*. Using a sensitive live-cell fluorescence assay we were able to benchmark at the single cell level IGR IRES translation efficiency against well characterized bacterial SD sequences *in vivo*. Using this quantitative approach, we were able to determine if translation of foreign mRNA is biased towards particular conserved structural features such of those present in IGR IRESs. Interestingly, disrupting the conserved PK structures (shown to be essential for IRES function in eukaryotes) does not always perturb IRES activity and instead often results in an increased translation efficiency. While the IRESs translation efficiency data do not align with a structure-driven interpretation, they strongly correlate with the mRNAs predicted free energy of folding (ρ = 0.61, Figure S10) and the translation strength predicted (Figure S11) using the Salis lab RBS calculator ^24^, suggesting that the PK structure in the IRES is in fact inhibitory to bacterial translation. Additionally, IGR IRES translation is strongly dependent on ribosomal protein S1 and the presence of a SD-like sequence upstream of the start codon. Collectively, our data demonstrates that the translational activity of RNAs derived from foreign DNA (foreign mRNAs) such as the IGR IRESs is not due to their three-dimensional structure and is primarily the result of the activity of ribosomal protein S1 and the overall robustness of canonical SD-dependent translation. It appears that in addition to its essential role facilitating efficient translation of endogenous mRNAs, S1 also enables the translation of structured foreign mRNAs, suggesting an important role for accessing genetic diversity trough horizontal gene transfer and therefore contributes to evolutionary fitness and stress response in bacteria.

While foreign mRNAs may have similar affinities to the prokaryotic ribosome/ribosomal subunits, demonstrating their ability to effectively compete for translation machinery, they still need to be transcribed at a sufficient quantity to be translated at functionally relevant levels. Although foreign DNA is not optimized for transcription in *E. coli*, sporadic transcription has been reported for AT-rich regions of DNA ^7^. In fact foreign DNA that transcribes too efficiently can be deleterious as it perturbs host transcription, for example, by sequestering RNA polymerase ^7,36,37^. Furthermore, foreign genes that are not AT-rich are less likely to be sporadically transcribed and therefore are unlikely to reach the proteome and to be kept by *E. coli*, evidenced by the fact that the genes ultimately retained have a relatively high AT-content ^38^. In the current study a T7 promoter/polymerase system was used for expression, resulting in transcription levels likely higher than for foreign DNA acquired by HGT. Using flow cytometry and RT-qPCR data we are able to compare the level of mRNA expression at our lowest observed fluorescence by using the reference gene *cysG*, which has been reported to be present in ∼2000 copies/0.1μg of total RNA ^39^. Just prior to induction the level of sfGFP mRNA is 3.3-fold higher than *cysG*, corresponding to roughly ∼6500 copies/0.1μg of total RNA. As *cysG* is a comparably low abundance mRNA ^39^ we can infer that our observations at least hold true for foreign mRNAs with transcription levels similar to *cysG*.

Once transcribed, mRNAs derived from the foreign genes must be translated to enter the proteome and provide their potential beneficial activity. To do so either the foreign mRNA will have to contain information (e.g., sequence, structure, etc.) allowing it to specifically interact with the bacterial translation machinery and circumvent the canonical bacterial translation initiation mechanism, or *E. coli* is inherently capable of translating foreign mRNAs by funneling them into the canonical mechanism. The latter is supported by recent findings that ribosomes with altered anti-SD sequences (incapable of base-pairing to canonical SDs) are able to initiate at the correct codons suggesting translation start sites are determined by inherent mRNA features such as upstream A-rich sequences and lower levels of surrounding mRNA structure ^29,40^. Interestingly, increased incidence of A-rich sequences is observed at almost every level (transcription, translation, gene retention) of foreign gene expression, suggesting the lower level of structure (DNA and RNA) is what allows these genes to be utilized. While foreign mRNAs may not have the same inherent features, incorrect start codon selection will only result in reduced translational efficiency of the respective mRNA. Most incorrect start sites will be out of frame and are unlikely to generate long or functional proteins. Those start sites that are in frame will either result in truncations or extensions of the protein both of which are likely to fold and retain at least some of the original enzymatic activity.

Interestingly in *E. coli, rpsA* (S1) shares an operon (and is co-transcribed) with *ihfB* which codes for a subunit of the host integration factor (Pedersen et al., 1984) a key element in HGT, gene expression, and DNA recombination. Data reported here supports a model where ribosomal protein S1 has a critical role in mediating the expression of foreign mRNAs (Figure 5). This is consistent with S1’s canonical role in host translation as it has already been shown to be essential for the translation of most endogenous *E. coli* mRNAs ^41^. During translation initiation S1 binds single stranded RNA in a sequence independent fashion, a critical role as many bacterial genes do not contain Shine-Dalgarno sequences ^33,41-44^. Additionally, S1 binds RNA containing pseudoknots and has been shown to be essential for docking and unfolding of structured mRNAs on the ribosome ^33,45^. Previous studies have reported that S1 can allow foreign mRNAs devoid of guanines (and therefore no SD sequence) from a plant infecting virus to form initiation complexes *in vitro* ^43^. It is tempting to imagine ribosomal protein S1 extending into solution and non-specifically binding mRNAs and “handing them over” to the ribosome increasing the local concentration in much the same manner as the ribosomal L7/L12 stalk for Elongation Factors Tu and G. Together with the *in vivo* data presented here, we propose that the bacterial translation machinery (like transcription) is fundamentally promiscuous as means of enabling the expression of foreign DNA acquired through HGT with S1 one playing a central role.

**Figure 5.**
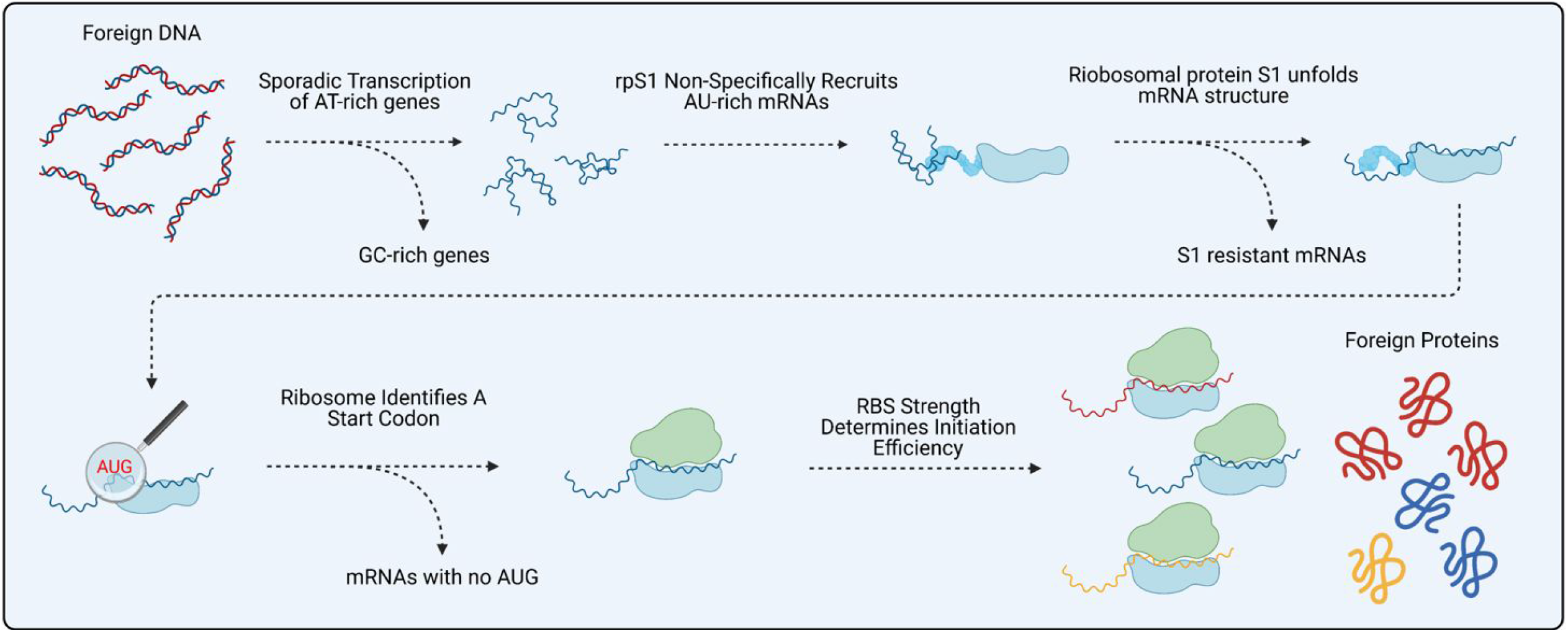
Proposed mechanism of S1 mediated expression of foreign genes. Foreign AT-rich DNA is sporadically transcribed before S1 non-specifically recruits it to the ribosome. If S1 can unfold the RNA structure and a start codon (AUG) can be identified translation occurs. The efficiency of this translation is dependent on the strength of the RBS (its ability to base pair to the anti-SDS).

Great care is taken by many organisms to control which genes are translated and to defend against expression of foreign DNA, as aberrant translation of proteins is implicated in several humans diseases ^46,47^. However, from the perspective of a bacterial population, the risks of sporadic deleterious gene expression by some of the bacteria are outweighed by the potential benefits making them more resilient to environmental stresses as the surviving bacterial cells will be able to repopulate. This strategy is commonly applied by bacteria to overcome environmental stress, for example by increasing the mutation rates to allow the survival of a portion of the population ^48^. On this background, uptake of genes that have already successfully evolved a specific stress-related beneficial function (e.g., antibiotic resistance genes) through HGT will provide an additional advantage, fast-tracking the adjustment of the bacterial population to the changing environmental stresses ^49^.

While the genes ultimately retained from horizontal gene transfer have biased biological function (e.g., antibiotic resistance), our data supports a model whereby the bacterial translation system has evolved to not be inherently biased for specific foreign mRNAs. Rather it allows foreign mRNA to be translated as long as the structure can be resolved (e.g., by ribosomal protein S1) upstream of an in frame start codon (Figure 5). Foreign mRNA even at lower transcription levels will be able to compete with endogenous mRNAs for translation by the ribosome primarily trough the strength of their “initiation region” upstream of the start codon, avoiding additional biases for expression, a leveling effect highlighting the central role of ribosomal protein S1 for bacterial physiology and fitness. With this in mind, the function of S1 during the expression of foreign mRNAs might be an interesting target for the development of therapeutic strategies addressing the spread of antimicrobial resistance by impairing the Horizontal Gene Transfer machinery of bacteria.

## ACKNOWLEDGEMENTS

We thank C. B. DeSouza, Q. Daviduck, Y. Yao, and S. Alvafor assistance with testing and data collection. We thank J. Heller for the 30S subunits lacking ribosomal protein S1. We thank the SynBridge synthetic biology maker space at the University of Lethbridge for access to scientific infrastructure.

## AUTHOR CONTRIBUTIONS

Conceptualization, L.R. and H.-J.W.; Methodology, L.R.; Investigation, L.R. and H.-J.W.; Writing - Original Draft, L.R.; Writing - Review & Editing, L.R. and H.-J.W.

## DECLERATION OF INTERESTS

The authors declare no competing interests.

## MATERIALS AND CORRESPONDANCE

For correspondence and materials requests contact H.-J. Wieden.

## METHODS

### Fluorescent reporter construct design

IRES reporter constructs were designed to adhere to BioBrick engineering standards^50^. BioBrick Prefix and Suffix sequences (RFC 10) flank each construct for ease of cloning into BioBrick vectors with standardized copy numbers. A T7 promoter (BBa_I719005) drives the transcription of optimized superfolder green fluorescent protein (sfGFP) coding sequence translationally controlled by an RBS or IRES sequences. Transcription is stopped by a transcriptional terminator (BBa_0015) downstream of the sfGFP coding sequence. Sequences for the strong (BBa_B0034), medium (BBa_B0032), and weak (BBa_B0033) RBSs were taken from the BioBrick part registry and the “dead” RBS is the reverse compliment of BBa_B0034. Sequences for CrPV (AF218039), IAPV (NC_009025.1), and PSIV (AB006531) IGR IRESs with 18 nts of corresponding downstream coding sequences were obtained from GenBank. We used the Salis Lab RBS calculator to ensure a strong RBS or an upstream start codon was not accidentally created during IRES mutagenesis and IRES scrambling^24^.

### Cloning and site directed mutagenesis

Fluorescent reporter constructs were synthesized (Integrated DNA Technologies and Twist Biosciences) and subcloned into pSB3C5, a medium to low copy number plasmid. Pseudoknot (PK) mutations and deletions were introduced using the Quickchange™ method. All reactions were carried out using a T_Gradient_ (Biometra) thermocycler and resulting mutant plasmids transformed into electro competent BL21-Gold (DE3) cells (Agilent). The integrity of all constructs and PK mutations were confirmed by sequencing (Genewiz).

### Cell Growth

50 mL of *E. coli* BL21 (Gold) DE3 cells containing fluorescent constructs were grown in LB media to mid log phase (0.5 OD_600nm_) at 37**°**C with shaking (200 rpm) in 125 mL Erlenmeyer flasks and expression induced with IPTG (1 mM final concentration). Cells were then harvested at distinct time intervals (fluorescent time courses) or grown for three hours before being analyzed by flow cytometry.

### Flow cytometry

Cells were pelleted, washed twice with and subsequently resuspended in FACSFlow™ (BD Biosciences), and kept on ice until cytometric analysis. Flow cytometry was performed on a BD FACSAria Fusion cell sorter (488 nm excitation, observing sfGFP fluorescence in the FITC channel) and data analysis performed on Flowjo software (Flowjo, LLC). All flow cytometry was performed in biological triplicate, collecting 100,000 events per replicate.

### sfGFP Immunoblotting

Whole cell lysate or 5 μL of PURExpress (New England BioLabs) reaction was loaded onto a nitrocellulose membrane (Pall Corporation) using a Biodot SF microfiltration apparatus (BioRad) and the presence of sfGFP was detected using an anti-GFP antibody (Abcam, ab6556) and a peroxidase conjugated secondary antibody (Sigma, A0545). Chemiluminescence from three biological replicates was quantified using an Amersham Imager 600 (GE healthcare).

As a secondary check regarding the effect the maturation time of the GFP has on signal generation, we used the GFP specific antibody to probe protein levels during the expression time course, which confirmed that our live cell fluorescence assay was accurately reporting protein levels and not variations in the sfGFP maturation times (Figure S4).

### RT-qPCR

Total RNA from three biological replicates was extracted from *E. coli* using an EZ-10 total RNA purification kit (Bio Basic) and the integrity/purity confirmed using formaldehyde agarose gel electrophoresis and A_260_/A_280_ ratio (Biodrop). Using 100 ng of total RNA and the respective reverse primers (IDT) (sfGFP 5’-GATAACGAGCAAAGCACTGAAC-3’ and cysG 5’-ATGCGGTGAACTGTGGAATAAACG-3’) cDNA was generated using qScript cDNA Supermix (Quanta Biosciences) according to manufacturer’s specifications. Quantitative PCR was performed according to manufacturer’s specifications on a StepOnePlus Real-Time PCR System (Thermo Fisher) using PerfeCTa® SYBR® Green SuperMix (Quantabio) with the corresponding forward primers (IDT) (sfGFP 5’-GGTGACGCAACTAATGGTAAAC-3’ and cysG 5’-TTGTCGGCGGTGGTGATGTC-3’) and the above reverse primers. All sfGFP mRNA threshold values were scaled relative to the accompanying cysG reference mRNA threshold values to account for differences in cDNA input.

### sfGFP Degradation Assay

50 mL of *E. coli* BL21 (Gold) DE3 cell containing fluorescent constructs were grown in LB media to mid log phase (0.6 OD_600nm_) at 37**°**C with shaking (200 rpm) in 125 mL Erlenmeyer flasks in the presence of IPTG (1 mM final concentration). 50 mL of cells were pelleted and washed twice in ABT minimal media (without IPTG) before being resuspended in 50 mL of ABT minimal media and incubated at 37**°**C with shaking (200 rpm) in 125 mL Erlenmeyer flasks ^51^. Cells were then harvested at specific time intervals and sfGFP measured by flow cytometry (as above). The OD_600nm_ was constant at ∼0.6 over the course of the experiment ensuring cells were not actively dividing. No variations in sfGFP levels or degradation rates (Figure S5) could be detected for the different constructs; in particular, the decay rate was so slow (sfGFP was stable over numerous days) that sfGFP degradation is negligible over the time of our experiments.

### RNA in vitro transcription, [^32^P] labelling, and purification

DNA templates for *in vitro* transcription were generated by PCR using plasmids containing either wild type or mutant IRESs. The obtained DNA was used in subsequent *in vitro* transcription reactions, and the resultant RNA purified by nucleic acid spin column (Bio Basic). The purity and homogeneity of the RNA was assessed by urea PAGE and A_260_/A_280_ ratio (BioDrop μLite, BioDrop).

500 ng of IRES RNA in water was unfolded by heating to 95°C for 2 min before being snap cooled on ice. RNA was then dephosphorylated by incubating at 37°C with Shrimp Alkaline Phosphatase (0.001 U/μL final concentration, Fermentas) for 60 min. Two hundred and fifty ng of dephosphorylated RNA were incubated with T4 polynucleotide kinase (0.5 U/μL final concentration, Fermentas) and 1.5 μL of [^32^P]-γ-ATP (30 μL total reaction volume) for 60 min at 37°C. To quench the reaction 1.5 μL of 0.5 M EDTA pH 8.0 was added and the reaction subsequently heated to 75°C for 10 min before the RNA was purified via EZ-10 Spin Column RNA Cleanup and Concentration Kit (Bio Basic).

### Purification of prokaryotic (70S) and eukaryotic (40S) ribosomes

Prokaryotic 30S ribosomal subunits were purified from *E. coli* MRE600 as per Becker *et al*.,^52,53^. Eukaryotic 40S ribosomal subunits were purified from HeLa cells (National Cell Culture Laboratory) as previously described ^54^.

### Removal of ribosomal protein S1 from 30S subunits (30S^*-S1*^ *subunits)*

30S ribosomal subunits were diluted tenfold in a high-salt dissociation buffer (20 mM Tris-HCl pH 7.5, 10 mM MgCl_2_, 60 mM KCl, 1 M NH_4_Cl, and 1 mM DTT). The mixture was incubated at 37 °C for 10 min before being added to poly(U) (Sigma Aldrich, P8563) and incubated at 4 °C for 1 hr with gentle inversion. The mixture was centrifuged at 500xg for 5 min, and the supernatant collected. The 30S^-S1^ subunits were pelleted via ultracentrifugation with a Sorvall S55-S swinging-bucket rotor ultracentrifuge (Thermo Scientific) at 55 000 rpm, at 4 °C for 24 hr and resuspended in TAKM_5_ (50 mM Tris-HCl pH 7.6, 70 mM NH_4_Cl, 30 mM KCl, 5 mM MgCl_2_) to a concentration of ∼15 μM. S1 removal was confirmed via SDS-PAGE and mass spectrometry (U of L Mass Spectrometry Facility).

### Nitrocellulose filtration assays

Radio-labeled RNA (50 nM final) in TAKM_5_ buffer was heated to 95°C for 10 min and slow cooled to room temperature. RNA was then incubated with increasing amounts of ribosomal subunits/ribosomes for 15 min at 37°C before being rapidly filtrated through a cellulose nitrate membrane filter (0.2μm, GE Healthcare). The cellulose nitrate membranes were washed with 1 mL of cold TAKM_5_ buffer and placed into 10 mL of EcoLite (+) scintillation cocktail (MP Bio), vortexed for 30 sec, and subsequently incubated at room temperature for 30 min, followed by vigorous mixing for 30 sec. The retained radioactivity was quantified by scintillation counting (Tri-carb 2810 TR LSA, Perkin Elmer). To ensure our system was able to replicate previously published data we also performed binding assays with HeLa 40S subunits. Consistent with previous work the WT CrPV IRES bound the 40S with a K_D_ of ∼14nM, while disruption of PK1 had no effect on 40S binding and disruption PK1 and PK3 in combination abolished 40S binding (Table S1)^12,55^.

### RNA Fluorescence Titration

Purified WT CrPV IGR IRES RNA was labelled at the 3’ end with pyrene as per Keffer-Wilkes *et al*.^56^. Labeled RNA (50 nM final) in TAKM_5_ was heated to 95°C for 10 min and slow cooled to room temperature. RNA was then incubated with increasing amounts of ribosomal subunits/ribosomes before being excited at 341 nm. The peak fluorescence at 391 nm was recorded and plotted as a function of increasing 30S concentration.

### Statistical Information

For all statistical analyses n=3 unless otherwise stated.

#### A) Flow Cytometry

Flow cytometry was performed as described in methods description. Mean fluorescence was calculated using FlowJo software. Standard deviation and relative significance (T-Test, two tailed) of each data set was calculated using Microsoft Excel.

#### B) Immunoblot

Immunoblot intensity was determined using the ImageJ gel analysis package. Mean intensity and standard deviation was calculated using Microsoft Excel.

#### C) RT-qPCR

Threshold levels were generated by the StepOnePlus Real-Time PCR System and mRNA levels were calculated using Microsoft Excel.

