## Supplemental Data for "Ribosomal protein S1 plays a critical role in horizontal gene transfer by mediating the expression of foreign mRNAs"

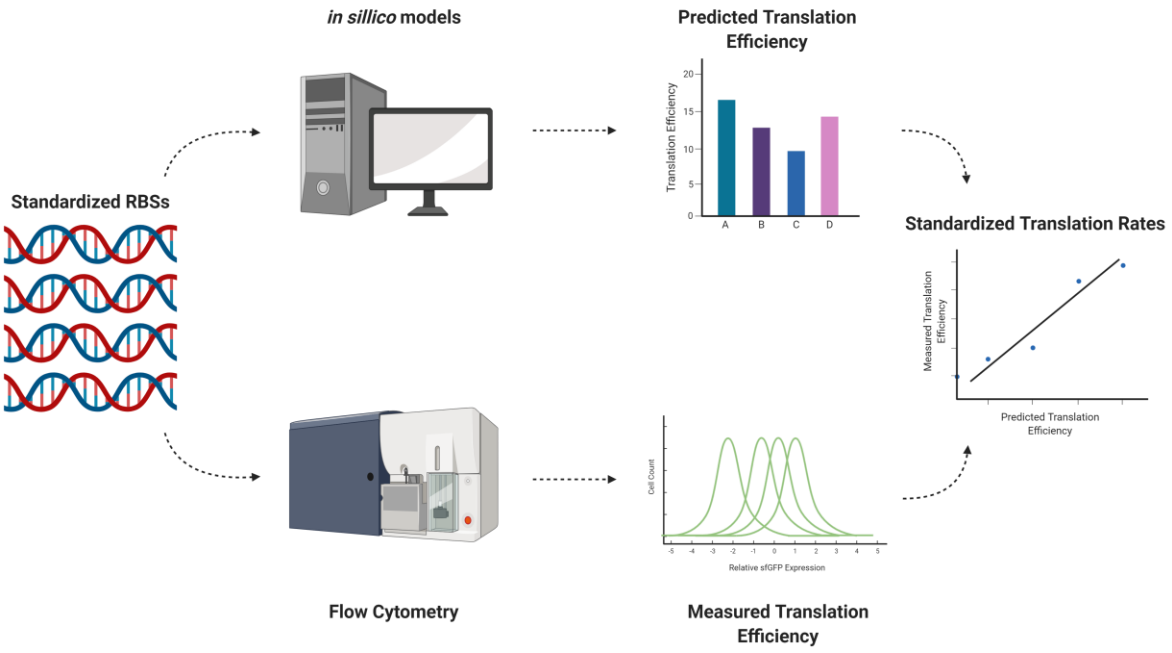


**Figure S1. Novel real-time fluorescence-based single-cell translation assay for benchmarking foreign gene expression *in vivo*.** Using standardized ribosome binding sites the translation efficiency of foreign translation elements can be benchmarked against canonical bacterial translation.

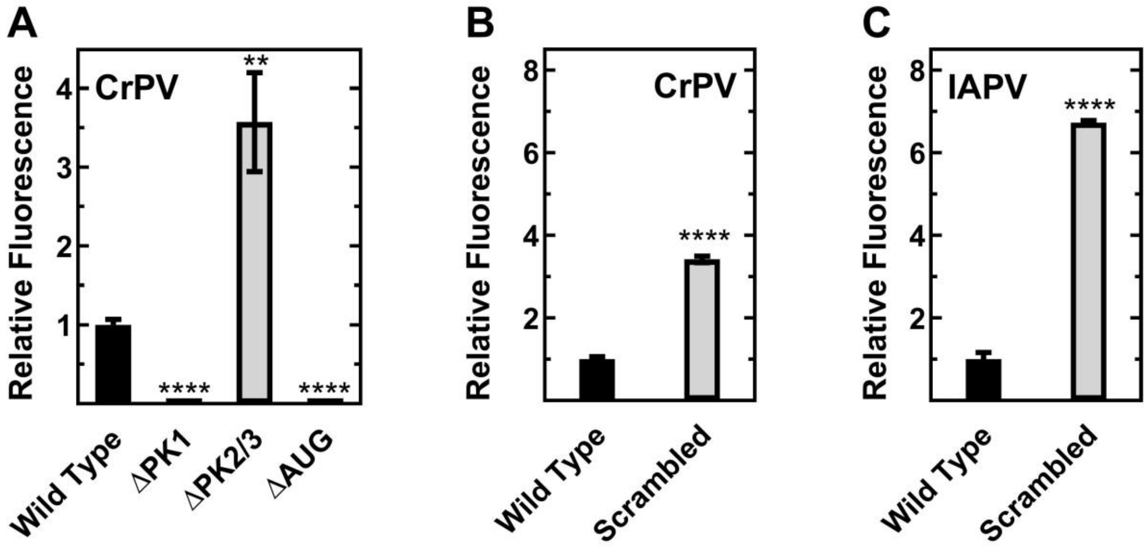


**Figure S2. Pseudoknot deletion and Scrambled IGR IRES variants have translation efficiency inconsistent with a structure based mechanism.** Relative fluorescence of *E. coli* containing IGR IRES constructs. Translation efficiency measured by flow cytometry, mean values of three biological replicates are plotted relative to the respective WT IRES; error bars indicate one standard deviation. Constructs with statistically significant differences from WT are indicated (* = P < 0.05, ** = P < 0.01, *** = P < 0.001, **** = P < 0.0001).

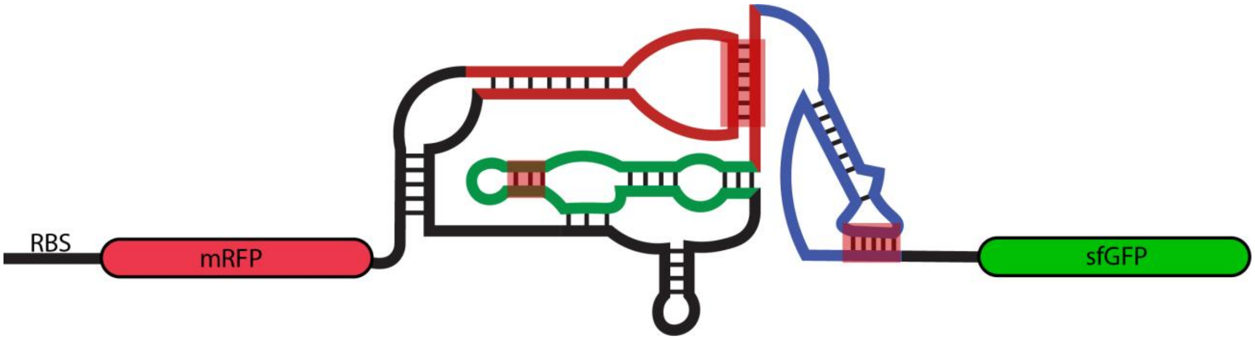


**Figure S3. Cartoon representation of the bicistronic fluorescent reporter construct including the secondary structure of the IGR IRESs.** Pseudoknots (PK) 1 (blue), 2 (red), and 3 (green) are indicated. Location of pseudoknot mutations are highlighted in red, the exact sequences are summarized in Table S4.

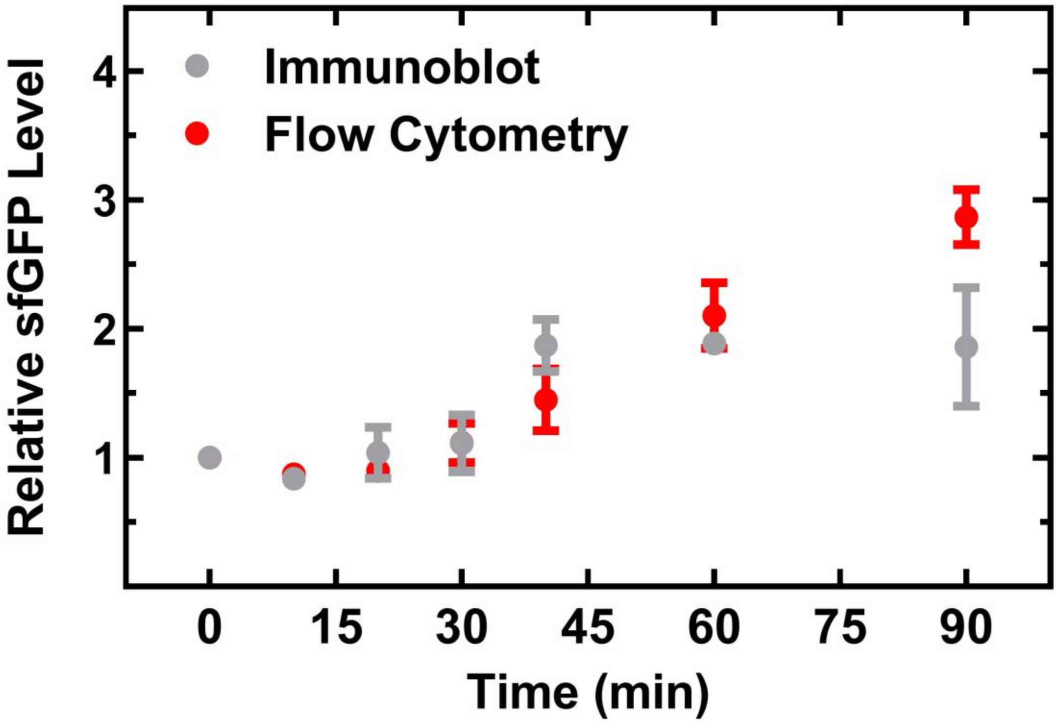


**Figure S4. Flow cytometry accurately reports the amount of sfGFP protein *in vivo*.** Relative fluorescence time course of *E. coli* containing PSIV IGR IRES constructs. Comparison between PK2_K/O flow cytometry data (fluorescence, red) and PK2_K/O immunoblot data (protein level, grey). Mean values of three biological replicates are plotted; error bars indicate one standard deviation.

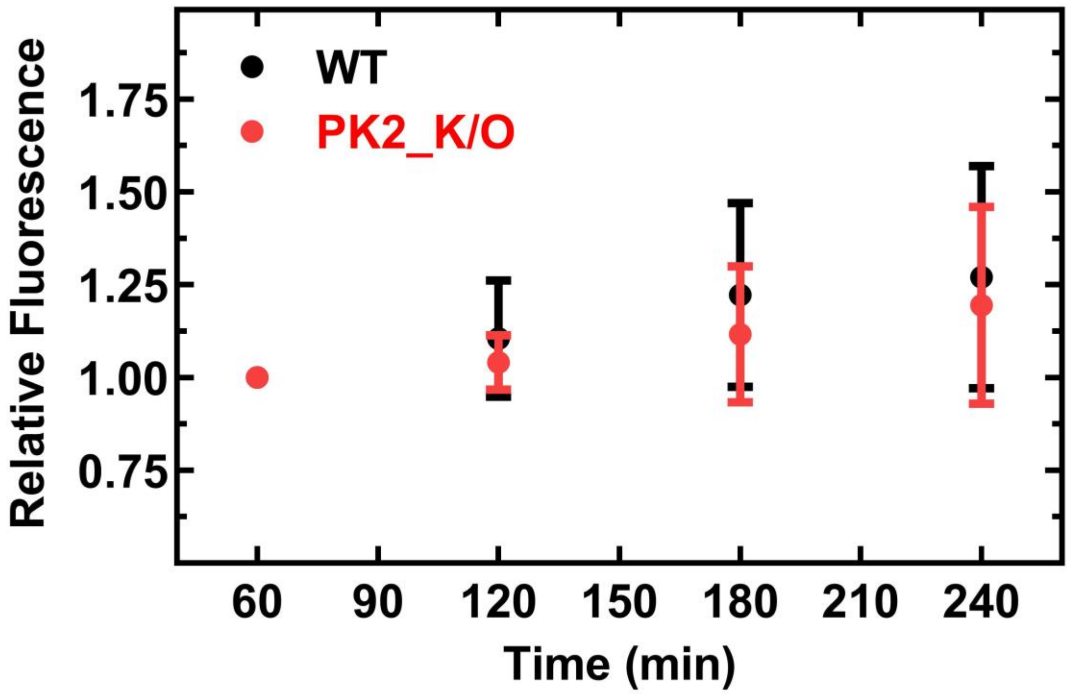


**Figure S5**. **Superfolder green fluorescent protein (sfGFP) is stable over multiple hours *in vivo***. Relative fluorescence time course of *E. coli* containing PSIV IGR IRES constructs post shift to minimal media measured by flow cytometry. Mean values of three biological replicates are plotted; error bars indicate one standard deviation.

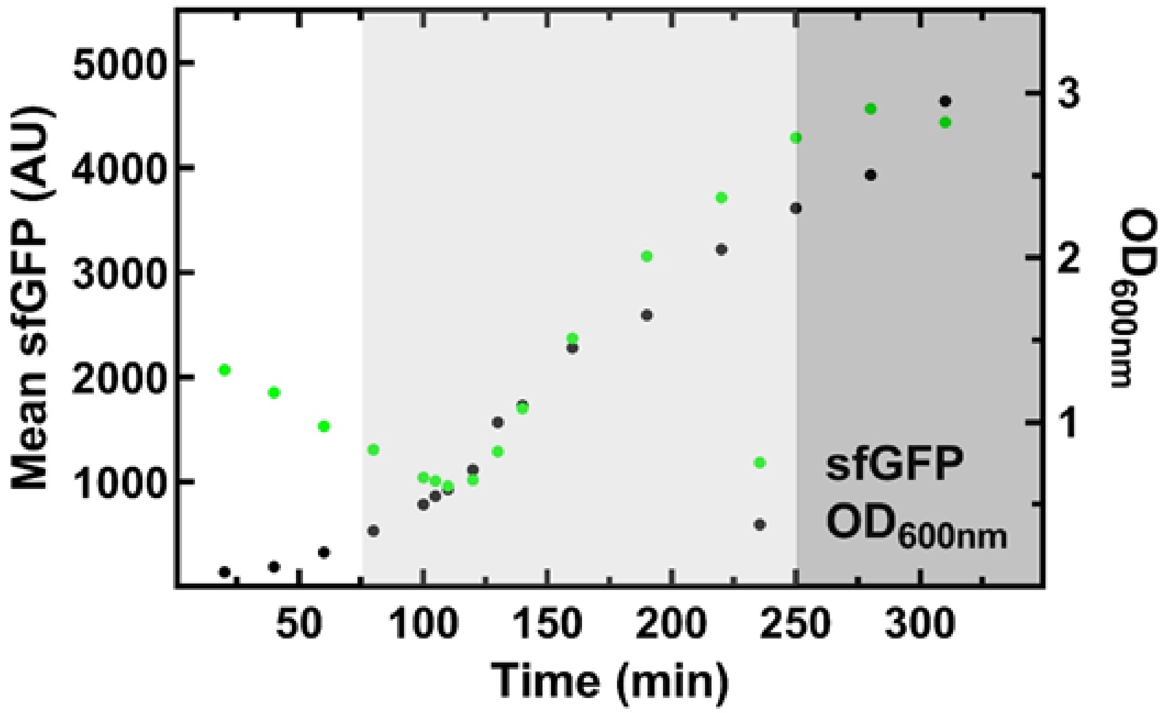


**Figure S6. Time course of sfGFP expression over several phases of cell growth.** Relative fluorescence *in vivo* time course (green) of *E. coli* containing the WT PSIV IGR IRES construct as measured by flow cytometry compared to the optical density at 600nm (black). Growth phases are indicated by the white (lag), light grey (exponential), and dark grey (early stationary) shading. Fluorescence decreased over the first 120 min as cells divide and dilute the sfGFP before expression is induced at 120min (0.6OD 600nm) with IPTG.

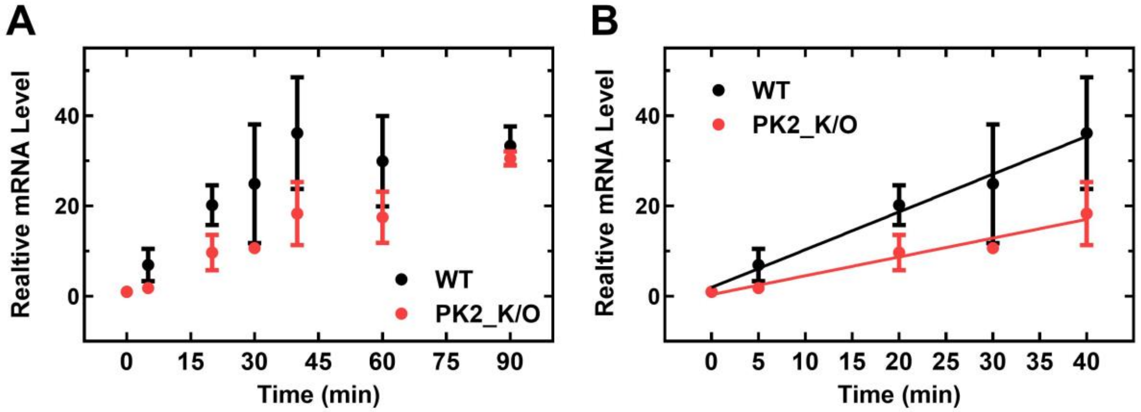


**Figure S7. PK2_K/O mRNA is less table than WT IRES mRNA *in vivo*.** sfGFP mRNA levels over several growth phases *in vivo*. (A) Relative mRNA level time course of live *E. coli* containing PSIV IGR IRES constructs measured by RT-qPCR. (B) Linear portion of mRNA expression from panel A. Mean values of three biological replicates are plotted; error bars indicate one standard deviation.

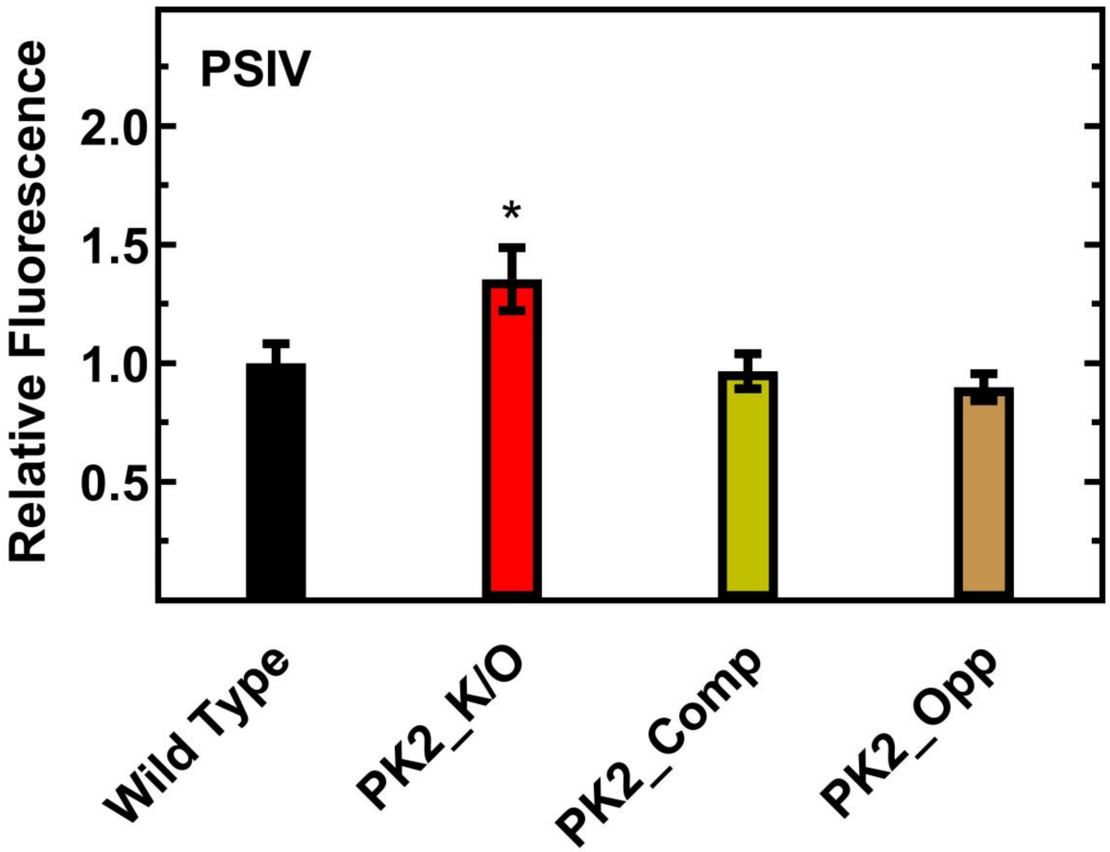


**Figure S8. Compensatory mutations restores PK2_K/O IRES translation efficiency to wild type**. Mean fluorescent values of three biological replicates obtained by flow cytometry are plotted relative to the respective WT IRES; error bars indicate one standard deviation. Constructs with statistically significant differences from WT are indicated (* = $P < 0.05$)

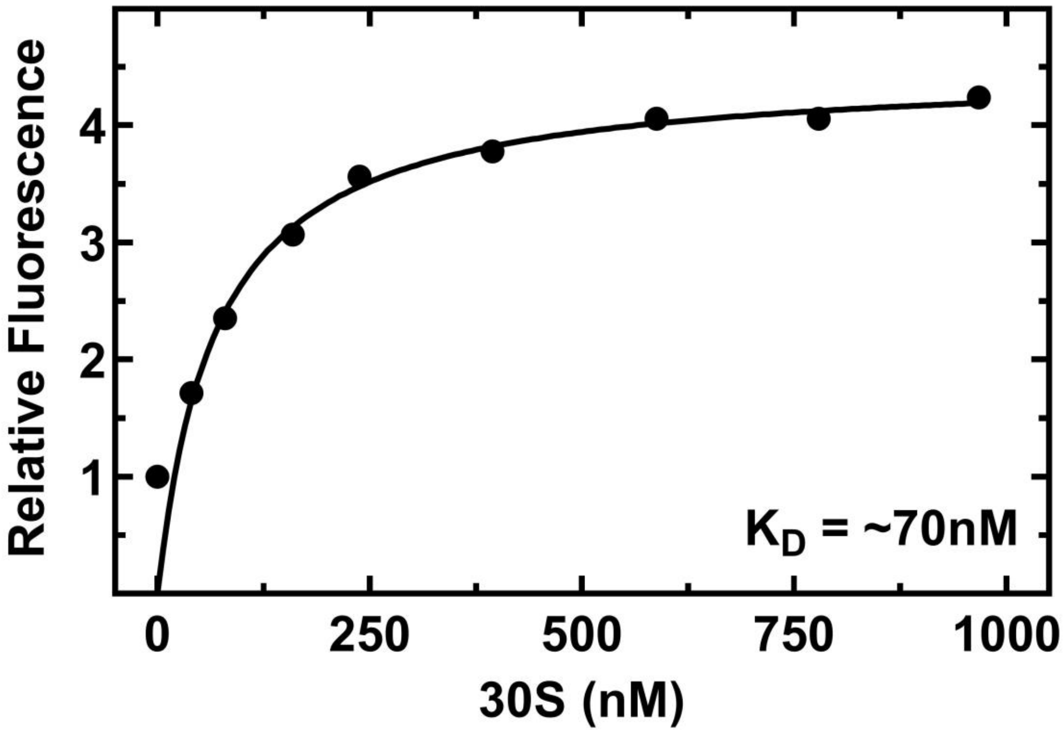


**Figure S9**. **The CrPV IGR IRES binds the 30S subunit with a nanomolar affinity.** Titration of fluorescently labelled WT CrPV IGR IRES with the 30S ribosomal subunit. Fluorescently labeled WT CrPV IGR IRES RNA is incubated with increasing amounts of 30S subunits. Relative fluorescence emission at 391 nm is shown ($\lambda_{ex}$ = 341 nm).

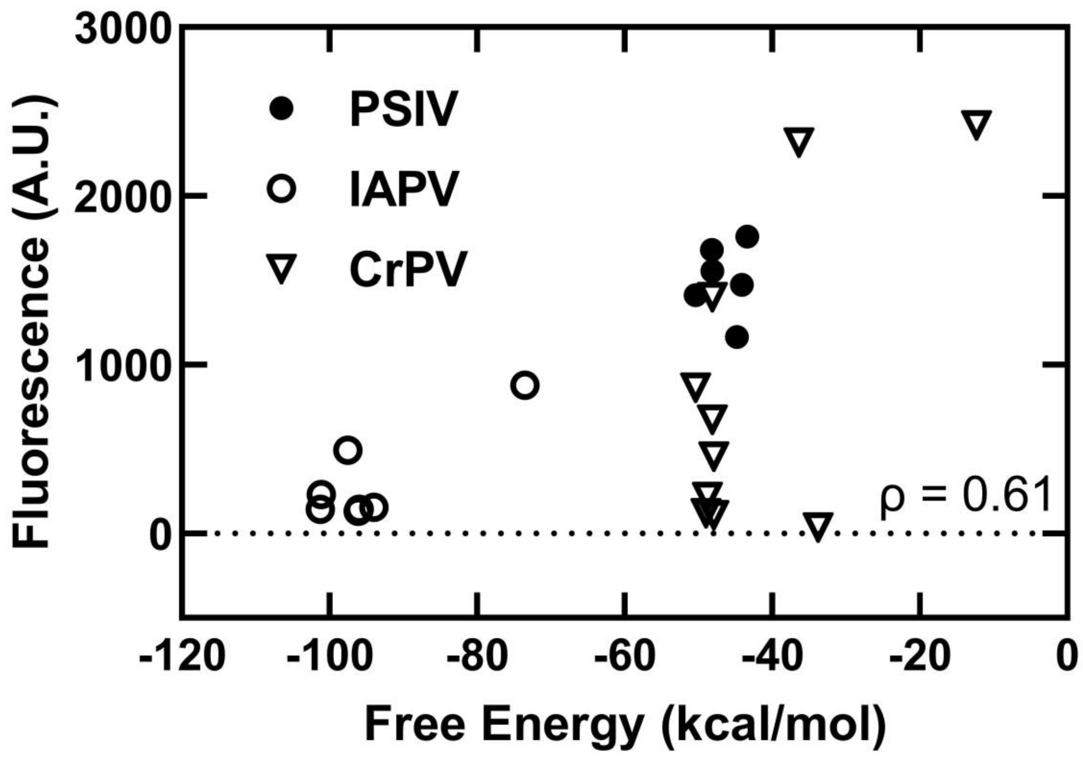


**Figure S10. Predicted free energy and translation efficiency of IGR IRES constructs are correlated.** Free energy predicted using mfold, translation efficiency measured by flow cytometry. Mean values of three biological replicates are plotted.

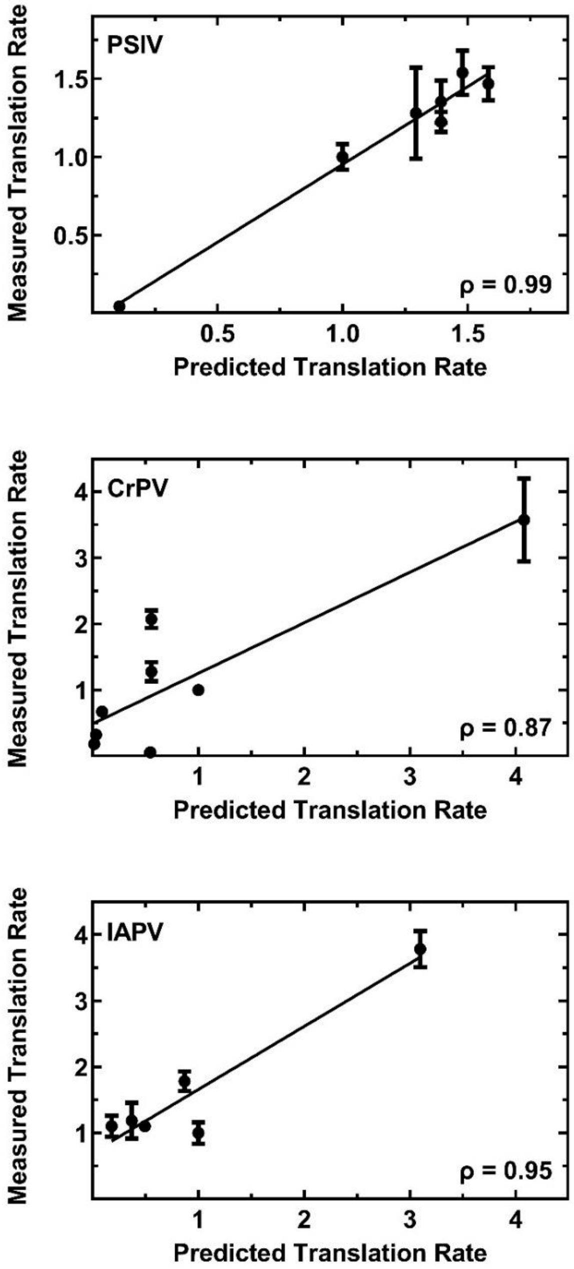


**Figure S11. Correlation of predicted and measured translation efficiency of IGR IRES constructs.** Translation efficiency predicted using the Salis lab RBS calculator, translation efficiency measured by flow cytometry. Mean values of three biological replicates are plotted; error bars indicate one standard deviation.

**Table S1.** Equilibrium dissociation constants and percent of RNA bound for CrPV IGR IRES variants and the 40S (HeLa) ribosomal subunit. Mean values of three replicates are plotted; error bars indicate one standard deviation.

| **CrPV Construct** | **$K_D$ (nM)** | **Percent Bound (%)** |
|---|---|---|
| Wild type | 14 ± 8 | 97 ± 7 |
| PK1_K/O | 4 ± 2 | 34 ± 2 |
| PK1+PK3_K/O | No Binding | No Binding |

**<u>Table S2.</u>** Equilibrium dissociation constants and percent of RNA bound for control RNAs and the 30S ribosome. Mean values of three replicates are plotted; error bars indicate one standard deviation.

| **Construct** | **$K_D$ (nM)** | **Percent Bound (%)** |
|---|---|---|
| rpsO | 10 ± 5 | 39 ± 2 |
| sodB | 13 ± 2 | 41 ± 3 |
| $tRNA^{Phe}$ | 35 ± 10 | 10 ± 1 |

**Table S3.** Equilibrium dissociation constants and percent of RNA bound for control RNAs and the 70S ribosome. Mean values of three replicates are plotted; error bars indicate one standard deviation.

| **Construct** | **$K_D$ (nM)** | **Percent Bound (%)** |
|---|---|---|
| rpsO | 94 ± 14 | 44 ± 10 |
| sodB | 47 ± 13 | 56 ± 8 |
| $tRNA^{Phe}$ | No Binding | No Binding |

**Table S4.** DNA sequences of monocistronic and bicistronic* constructs.

| CrPV | Sequence Upstream from sfGFP |
|---|---|
| WT | 5'-<br>AAAGCAAAAATGTGATCTTGCTTGTAAATACAATTTTGAGAGGTTAAT<br>AAATTACAAGTAGTGCTATTTTTGTATTTAGGTTAGCTATTTAGCTTTA<br>CGTTCCAGGATGCCTAGTGGCAGCCCCACAATATCCAGGAAGCCCT<br>CTCTGCGGTTTTTCAGATTAGGTAGTCGAAAAACCTAAGAAATTTAC<br>CTGCTACATTTCAAGATAAA-3' |
| PK1_K/O | 5'-<br>AAAGCAAAAATGTGATCTTGCTTGTAAATACAATTTTGAGAGGTTAAT<br>AAATTACAAGTAGTGCTATTTTTGTATTTAGGTTAGCTATTTAGCTTTA<br>CGTTCCAGGATGCCTAGTGGCAGCCCCACAATATCCAGGAAGCCCT<br>CTCTGCGGTTTTTCAGATTAGGTAGTCGAAAAACCTAAGAAATTTAG<br>GTGCTACATTTCAAGATAAA-3' |
| PK2_K/O | 5'-<br>AAAGCAAAAATGTGATCTTGCTTGTAAATACAATTTTGAGAGGTTAAT<br>AAATTACAAGTAGTGCTATTTTTGTATTTAGGTTAGCTATTTAGCTTTA<br>CGTTCCAGGATGCCTAGTGGCAGCCCCACAATATCCAGGAAGCGGA<br>GAGTGCGGTTTTTCAGATTAGGTAGTCGAAAAACCTAAGAAATTTAC<br>CTGCTACATTTCAAGATAAA-3' |
| PK3_K/O | 5'-<br>AAAGCAAAAATGTGATCTTGCTTGTAAATACAATTTTGAGAGGTTAAT<br>AAATTACAAGTAGTGCTATTTTTGTATTTAGGTTAGCTATTTAGCTTTA |

| | |
|---|---|
| | CGTTCCAGGATGCCTAGTGGCAGCCCGTGAATATCCAGGAAGCCCT CTCTGCGGTTTTTCAGATTAGGTAGTCGAAAAACCTAAGAAATTTAC CTGCTACATTTCAAGATAAA-3' |
| PK1+PK3_ K/O | 5'- AAAGCAAAAATGTGATCTTGCTTGTAAATACAATTTTGAGAGGTTAAT AAATTACAAGTAGTGCTATTTTTGTATTTAGGTTAGCTATTTAGCTTTA CGTTCCAGGATGCCTAGTGGCAGCCGGTGAATATCCAGGAAGCCCT CTCTGCGGTTTTTCAGATTAGGTAGTCGAAAAACCTAAGAAATTTAG GTGCTACATTTCAAGATAAA |
| PK1+PK2+ PK3_K/O | 5'- AAAGCAAAAATGTGATCTTGCTTGTAAATACAATTTTGAGAGGTTAAT AAATTACAAGTAGTGCTATTTTTGTATTTAGGTTAGCTATTTAGCTTTA CGTTCCAGGATGCCTAGTGGCAGCCGGTGAATATCCAGGAAGCGGA GAGTGCGGTTTTTCAGATTAGGTAGTCGAAAAACCTAAGAAATTTAG GTGCTACATTTCAAGATAAA-3' |
| PK1 Deletion | 5'- AAAGCAAAAATGTGATCTTGCTTGTAAATACAATTTTGAGAGGTTAAT AAATTACAAGTAGTGCTATTTTTGTATTTAGGTTAGCTATTTAGCTTTA CGTTCCAGGATGCCTAGTGGCAGCCCCACAATATCCAGGAAGCCCT CTCTGCCTACATTTCAAGATAAA-3' |
| PK2+PK3 Deletion | 5'- GCGGTTTTTCAGATTAGGTAGTCGAAAAACCTAAGAAATTTACCTGC TACATTTCAAGATAAA-3' |

| | |
|---|---|
| Scrambled | 5'-<br>GTAAGATGTTTCATGACCAACCCTAAATCTAAGTATATAAGTTGCACT<br>TGAAAATGGTAATTCTTAATAAGCTGGATGAGTAACGAGTATTATTAT<br>CGAACGGAATTTTCCTGGTGAAACACTCTTTCATGACACGGAAATAA<br>GGTGCGTCTTCTTTCAATATATCTCACGATGGCTTGAACAGACGGTT<br>TCATAACTTTCTTAAATGCG-3' |
| IAPV | Sequence Upstream from sfGFP |
| WT | 5'-<br>GAGCGGTTTCTGGAATACTATATGTAAGTATAGTGTTCTGGAGGCAT<br>CATTCTATGGTTACCCATCATTAGAGGAAATTTCCAATAAACTCTGGT<br>GTAAGGCTTAGAGTGATGGTCGAGGTGCCCTATTTAGGGTGAGGAG<br>CCTCGGTGGCAGCCCCACCAAATCCTCTATTGGATAGGAACAGCTG<br>TACTGGGCAGTTACAGCAGTCGTATGGTAACACATGCGGCGTTCCG<br>AAATACCATGCCTGGCGATTCACAACAAGAA-3' |
| PK1_K/O | 5'-<br>GAGCGGTTTCTGGAATACTATATGTAAGTATAGTGTTCTGGAGGCAT<br>CATTCTATGGTTACCCATCATTAGAGGAAATTTCCAATAAACTCTGGT<br>GTAAGGCTTAGAGTGATGGTCGAGGTGCCCTATTTAGGGTGAGGAG<br>CCTCGGTGGCAGCCCCACCAAATCCTCTATTGGATAGGAACAGCTG<br>TACTGGGCAGTTACAGCAGTCGTATGGTAACACATGCGGCGTTCCG<br>AAATACCATGGGTGGCGATTCACAACAAGAA-3' |
| PK2_K/O | 5'-<br>GAGCGGTTTCTGGAATACTATATGTAAGTATAGTGTTCTGGAGGCAT |

| | |
|---|---|
| | CATTCTATGGTTACCCATCATTAGAGGAAATTTCCAATAAACTCTGGT GTAAGGCTTAGAGTGATGGTCGAGGTGCCCTATTTAGGGTGAGGAG CCTCGGTGGCAGCCCCACCAAATCCTCTTAACCATAGGAACAGCTG TACTGGGCAGTTACAGCAGTCGTATGGTAACACATGCGGCGTTCCG AAATACCATGCCTGGCGATTCACAACAAGAA-3' |
| PK3_K/O | 5'- GAGCGGTTTCTGGAATACTATATGTAAGTATAGTGTTCTGGAGGCAT CATTCTATGGTTACCCATCATTAGAGGAAATTTCCAATAAACTCTGGT GTAAGGCTTAGAGTGATGGTCGAGGTGCCCTATTTAGGGTGAGGAG CCTCGGTGGCAGCCCCTGGAAATCCTCTATTGGATAGGAACAGCTG TACTGGGCAGTTACAGCAGTCGTATGGTAACACATGCGGCGTTCCG AAATACCATGCCTGGCGATTCACAACAAGAA-3' |
| PK1+PK3_ K/O | 5'- GAGCGGTTTCTGGAATACTATATGTAAGTATAGTGTTCTGGAGGCAT CATTCTATGGTTACCCATCATTAGAGGAAATTTCCAATAAACTCTGGT GTAAGGCTTAGAGTGATGGTCGAGGTGCCCTATTTAGGGTGAGGAG CCTCGGTGGCAGCCCCTGGAAATCCTCTATTGGATAGGAACAGCTG TACTGGGCAGTTACAGCAGTCGTATGGTAACACATGCGGCGTTCCG AAATACCATGGGTGGCGATTCACAACAAGAA-3' |
| PK1+PK2+ PK3_K/O | 5'- GAGCGGTTTCTGGAATACTATATGTAAGTATAGTGTTCTGGAGGCAT CATTCTATGGTTACCCATCATTAGAGGAAATTTCCAATAAACTCTGGT GTAAGGCTTAGAGTGATGGTCGAGGTGCCCTATTTAGGGTGAGGAG |

| | |
|---|---|
| | CCTCGGTGGCAGCCCCTGGAAATCCTCTTAACCATAGGAACAGCTG TACTGGGCAGTTACAGCAGTCGTATGGTAACACATGCGGCGTTCCG AAATACCATGGGTGGCGATTCACAACAAGAA-3' |
| Scrambled | 5'- CCCGTATGGGGATCGGACCGTGTTGCGCGCGATTGGATATACAAGC ATAATTCTAAAAGGGACCTTGTCTGGGTTATCACTCCGATCTTGCGT TAACCATTATGTTATAACGCAGACTATTGAGCCGCGAGACAAAGCCC CTTTGATTTTAAGTGACATCACTAGGTCAAGCAGCAAGGTCTGGCAG ACTAGGTATTAAGTAAAGTGTGCATACGGTGAGTGTTGAGGCGCCG CCAGTCAGCGGGATAAAAGATCTTAGCTTAT-3' |
| PSIV | Sequence Upstream from sfGFP |
| WT | 5'- AAGCTGACTATGTGATCTTATTAAAATTAGGTTAAATTTCGAGGTTAA AAATAGTTTTAATATTGCTATAGTCTTAGAGGTCTTGTATATTTATACT TACCACACAAGATGGACCGGAGCAGCCCTCCAATATCTAGTGTACC CTCGTGCTCGCTCAAACATTAAGTGGTGTTGTGCGAAAAGAATCTCA CTTCAAGAAAAAGAATTTACC-3' |
| PK1_K/O | AAGCTGACTATGTGATCTTATTAAAATTAGGTTAAATTTCGAGGTTAA AAATAGTTTTAATATTGCTATAGTCTTAGAGGTCTTGTATATTTATACT TACCACACAAGATGGACCGGAGCAGCCCTCCAATATCTAGTGTACC CTCGTGCTCGCTCAAACATTTTCACGTGTTGTGCGAAAAGAATCTCA CTTCAAGAAAAAGAATTTACC-3' |

| | |
|---|---|
| PK2_K/O | 5'- AAGCTGACTATGTGATCTTATTAAAATTAGGTTAAATTTCGAGGTTAA AAATAGTTTTAATATTGCTATAGTCTTAGAGGTCTTGTATATTTATACT TACCACACAAGATGGACCGGAGCAGCCCTCCAATATCTAGTGTACG GAGCTGCTCGCTCAAACATTAAGTGGTGTTGTGCGAAAAGAATCTCA CTTCAAGAAAAAGAATTTACC-3' |
| PK2_Opp | 5'- AAGCTGACTATGTGATCTTATTAAAATTAGGTTAAATTTGCTCCTTAA AAATAGTTTTAATATTGCTATAGTCTTAGAGGTCTTGTATATTTATACT TACCACACAAGATGGACCGGAGCAGCCCTCCAATATCTAGTGTACC CTCGTGCTCGCTCAAACATTAAGTGGTGTTGTGCGAAAAGAATCTCA CTTCAAGAAAAAGAATTTACC-3' |
| PK2_Comp | 5'- AAGCTGACTATGTGATCTTATTAAAATTAGGTTAAATTTGCTCCTTAA AAATAGTTTTAATATTGCTATAGTCTTAGAGGTCTTGTATATTTATACT TACCACACAAGATGGACCGGAGCAGCCCTCCAATATCTAGTGTACG GAGCTGCTCGCTCAAACATTAAGTGGTGTTGTGCGAAAAGAATCTCA CTTCAAGAAAAAGAATTTACC-3' |
| PK3_K/O | 5'- AAGCTGACTATGTGATCTTATTAAAATTAGGTTAAATTTCGAGGTTAA AAATAGTTTTAATATTGCTATAGTCTTAGAGGTCTTGTATATTTATACT TACCACACAAGATGGACCGGAGCAGCCGAGGAATATCTAGTGTACC |

| | |
|---|---|
| | CTCGTGCTCGCTCAAACATTAAGTGGTGTTGTGCGAAAAGAATCTCA CTTCAAGAAAAAGAATTTACC-3' |
| PK1+PK3_ K/O | 5'- AAGCTGACTATGTGATCTTATTAAAATTAGGTTAAATTTCGAGGTTAA AAATAGTTTTAATATTGCTATAGTCTTAGAGGTCTTGTATATTTATACT TACCACACAAGATGGACCGGAGCAGCCGAGGAATATCTAGTGTACC CTCGTGCTCGCTCAAACATTTTCACGTGTTGTGCGAAAAGAATCTCA CTTCAAGAAAAAGAATTTACC-3' |
| PK1+PK2+ PK3_K/O | 5'- AAGCTGACTATGTGATCTTATTAAAATTAGGTTAAATTTCGAGGTTAA AAATAGTTTTAATATTGCTATAGTCTTAGAGGTCTTGTATATTTATACT TACCACACAAGATGGACCGGAGCAGCCGAGGAATATCTAGTGTACG GAGCTGCTCGCTCAAACATTTTCACGTGTTGTGCGAAAAGAATCTCA CTTCAAGAAAAAGAATTTACC-3' |
| SDS- like_K/O | 5'- AAGCTGACTATGTGATCTTATTAAAATTAGGTTAAATTTCGAGGTTAA AAATAGTTTTAATATTGCTATAGTCTTAGAGGTCTTGTATATTTATACT TACCACACAAGATGGACCGGAGCAGCCCTCCAATATCTAGTGTACC CTCGTGCTCGCTCAAACATTAAGTGGTGTTGTGCGAAAAGAATCTCA CTTCAACTGGTCTTGACGGCC-3' |
| RBS | Sequence Upstream from sfGFP |

| | |
|---|---|
| Strong (BBa_B0034) | 5'-AAAGAGGAGAAATACTAG-3' |
| Medium (BBa_B0032) | 5'-TCACACAGGAAAGTACTAG-3' |
| Weak (BBa_B0033) | 5'-TCACACAGGACTACTAG-3' |
| Dead (BBa-B0034 Inv) | 5'-TTTCTCCTCTTTACTAG-3' |

*All PSIV bicistronic constructs are identical to the monocistronic constructs except they have the following sequence (coding for T7-RBS-mRFP) upstream of the IGR IRES: 5'-

TAATACGACTCACTATAGGGAGATACTAGAGTCACACAGGACTACTAGATGGCTTC
CTCCGAAGACGTTATCAAAGAGTTCATGCGTTTCAAAGTTCGTATGGAAGGTTCCG
TTAACGGTCACGAGTTCGAAATCGAAGGTGAAGGTGAAGGTCGTCCGTACGAAGG
TACCCAGACCGCTAAACTGAAAGTTACCAAAGGTGGTCCGCTGCCGTTCGCTTGG
GACATCCTGTCCCCGCAGTTCCAGTACGGTTCCAAAGCTTACGTTAAACACCCGG
CTGACATCCCGGACTACCTGAAACTGTCCTTCCCGGAAGGTTTCAAATGGGAACGT
GTTATGAACTTCGAAGACGGTGGTGTTGTTACCGTTACCCAGGACTCCTCCCTGCA
AGACGGTGAGTTCATCTACAAAGTTAAACTGCGTGGTACCAACTTCCCGTCCGACG
GTCCGGTTATGCAGAAAAAAACCATGGGTTGGGAAGCTTCCACCGAACGTATGTA

CCCGGAAGACGGTGCTCTGAAAGGTGAAATCAAAATGCGTCTGAAACTGAAAGAC

GGTGGTCACTACGACGCTGAAGTTAAAACCACCTACATGGCTAAAAAACCGGTTCA

GCTGCCGGGTGCTTACAAAACCGACATCAAACTGGACATCACCTCCCACAACGAA

GACTACACCATCGTTGAACAGTACGAACGTGCTGAAGGTCGTCACTCCACCGGTG

CTTAATAACGCTGATAGTGCTAGTGTAGATCGCTACTAGAG-3’

**Table S5.** DNA sequences for control RNAs used in filter binding.

| **Control RNAs** | **Sequences used in filter binding** |
|---|---|
| *rpsO* | 5'-GGATCCTAATACGACTCACTATAGGTACGAGTAGAATACTGCCGCTTAACGTCGCGTAAATTGTTTAACACTTTGCGTAACGTACACTGGGATCGCTGAATTAGAGATCGGCGTCCTTTCATTCTATATACTTTGGAGTTTTAAAATGTCTCAAGTACTGAAGCAACAGCTAAAATCGTTTCTGAGTTTGGTCGTGACGCAAACGACACC-3' |
| *sodB* | 5'-GGATCCTAATACGACTCACTATAGGATACGCACAATAAGGCTATTGTACGTATGCAAATTAATAATAAAGGAGAGTAGCAATGTCATTCGAATTACCTGCACTACCATATGCTAAAGATGCTCTGGCACCGCACATTTCTGCGG-3' |
| $tRNA^{Phe}$ | 5'-GCGCGGAUGCUCAGUCGGUAGAGCAGGGGAUUGAAAAUCCCCGUGUCCUUGGUUCGAUUCCGAGUCCGCGCACCA-3' |
